# Plant genetic effects on microbial hubs impact fitness across field trials

**DOI:** 10.1101/181198

**Authors:** Benjamin Brachi, Daniele Filiault, Hannah Whitehurst, Paul Darme, Pierre Le Gars, Marine Le Mentec, Timothy C. Morton, Envel Kerdaffrec, Fernando Rabanal, Alison Anastasio, Mathew S. Box, Susan Duncan, Feng Huang, Riley Leff, Polina Novikova, Matthew Perisin, Takashi Tsuchimatsu, Roderick Woolley, Caroline Dean, Magnus Nordborg, Svante Holm, Joy Bergelson

## Abstract

Although complex interactions between hosts and microbial associates are increasingly well documented, we still know little about how and why hosts shape microbial communities in nature. In addition, host genetic effects on microbial communities vary widely depending on the environment, obscuring conclusions about which microbes are impacted and which plant functions are important. We characterized the leaf microbiota of 200 *A. thaliana* genotypes in eight field experiments and detected consistent host effects on specific, broadly distributed microbial OTU’s. Host genetics disproportionately influenced hubs within the microbial communities, with their impact then percolating through the community, as evidenced by a decline in the heritability of particular OTUs with their distance to the nearest hub. By simultaneously measuring host performance, we found that host genetics associated with microbial hubs explained over 10% of the variation in lifetime seed production among host genotypes across sites and years. We successfully cultured one of these microbial hubs and demonstrated its growth-promoting effects on plants grown in sterile conditions. Finally, genome-wide association mapping identified many putatively causal genes with small effects on the relative abundance of microbial hubs across sites and years, and these genes were enriched for those involved in the synthesis of specialized metabolites, auxins and the immune system. Using untargeted metabolomics, we corroborate the consistent association of variation in specialized metabolites and microbial hubs across field sites. Together, our results reveal that host natural variation impacts the microbial communities in consistent ways across environments and that these effects contribute to fitness variation among host genotypes.

## Main

Hosts harbor complex microbial communities that are thought to impact health and development [1]. Human microbiota has been implicated in a variety of diseases, including obesity and cancer [2]. Efforts are thus underway to determine the host factors shaping these communities [3,4], and to use next-generation probiotics to inhibit colonization by pathogens [5]. Similarly, in agriculture, there is great hope of shaping the composition of the microbiota in order to mitigate disease and increase crop yield in a sustainable fashion. Indeed, the Food and Agriculture Organization of the United Nations has made the use of biological control and growth-promoting microbial associations a clear priority for improving food production [6].

Plant-associated microbes can be beneficial in many ways, including improving access to nutrients, activating or priming the immune system, and competing with pathogens. For example, seeds inoculated with a combination of naturally occurring microbes were found to be protected from a sudden-wilt disease that emerged after continuous cropping [7]. Thus, it would be advantageous to breed crops that promote the growth of beneficial microbes under a variety of field conditions, a prospect that is made more likely by the demonstration of host genotypic effects on their microbiota [8–10]. However, microbial communities are complex entities that are influenced by the combined impact of host factors, environment and microbe-microbe interactions [11]. Indeed, several studies have found a strong influence of the environment on estimates of host genotype effects [8,12,13]. Although most, if not all, studies exploring the influence that host genotype exerts on microbial communities suggest that such plant control could be beneficial to plant performance, almost nothing is known about the relationship between host genotype effects on microbial communities and on plant performance or fitness. As a consequence, the extent to which host plants can control microbial communities to their advantage, especially in a consistent manner across multiple environments, remains unclear.

Here, we combine large-scale field experiments in natural environments, extensive microbial community analysis, and genome-wide association mapping to: (i) determine how host genotype affects different microbial community members, and thus shapes the overall microbiome; (ii) estimate host genotype effects on microbial communities across eight environments and investigate the contribution of those effects to the performance of plant genotypes; and (iii) use genome-wide association mapping to identify key pathways that shape the leaf microbial communities across multiple environmental conditions.

### Snapshot of microbial community variation

We performed a set of field experiments that included natural inbred lines of *Arabidopsis thaliana* (hereafter “accessions”) originally collected throughout Sweden, mainly in two climatically contrasted regions of the country (Supplementary Table 1); *A. thaliana* in the north of Sweden experiences long, snowy winters, and as a consequence plants are typically found on south-facing slopes of rocky cliffs. *Arabidopsis* populations in the south of Sweden, on the other hand, tend to be associated with agricultural or disturbed fields that experience highly variable snow cover over the winter months. We used replicate experiments in four representative *Arabidopsis* sites, two each in the north (sites NM and NA) and south (sites SU and SR) of Sweden. Experiments were repeated across two years, for a total of eight experiments.

Each experiment was organized in a complete randomized block design including 24 replicates of 200 sequenced accessions [14], established as seedlings in a mixture of native and potting soil and timed to coincide with local germination flushes in late summer. Immediately upon snowmelt in early spring, we sampled and freeze-dried 5 to 6 whole rosettes per accession. DNA was extracted from the freeze-dried rosettes and both the ITS1 portion of the Internal Transcribed Spacer (ITS) and the V5 to V7 regions of the 16S RNA gene were sequenced to characterize the fungal and bacterial communities, respectively [9,11,15]. The sequences obtained were clustered into Operational Taxonomic Units (OTUs) using Swarm to generate community matrices [14] (see “Count table filtering” section in the methods). The frequency distributions of OTUs were highly skewed, with the top ten most common OTUs contributing on average 59% of the reads in each experiment (ranging from 45 to 78%). Taxonomic assignments indicate that the fungal communities were dominated by Leotimycetes and Dothideomycetes while the bacterial communities included high proportions of Alphaproteobacteria and Actinobacteria (Extended Data Fig. 1).

**Fig. 1.**
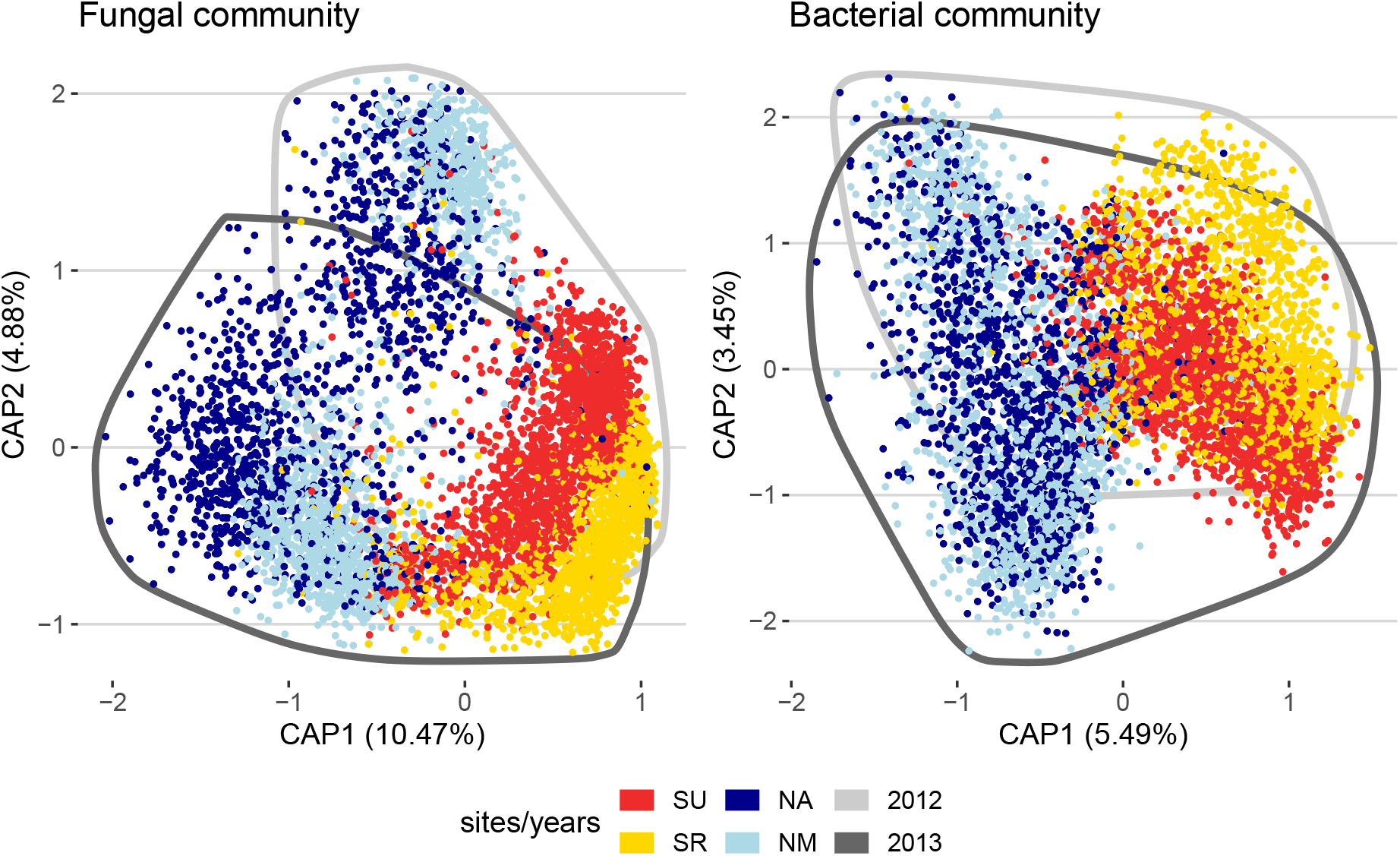
Plants grown in different environments have different microbial communities. The plots represent the projection of each sample on the plane defined by the first two constrained components of the fungal and bacterial communities, describing variation among sites and years. The percentages in parentheses are the proportion of the total inertia (square root of the Bray-Curtis dissimilarity) explained by each component. The colors of the points indicate the site from which samples were collected. Experiments from the South are represented in red (SU) and yellow (SR), and experiments from the North in light blue (NR) and dark blue (NA). All points from 2012 and 2013 are encircled by a dark and lighter grey line respectively.

In a principal coordinate analysis, differences between northern and southern sites explained 10 and 5% of the overall diversity in the fungal and bacterial communities, respectively, while differences between the two consecutive years explained 5 and 3%. This level of differentiation among experiments likely underestimates that present in the native soil, as it has been shown that hosts filter the microbial community to reduce site-to-site differences [17,18] (Fig. 1). In addition, there may have been a homogenizing effect of using a combination of local and potting soil. Irrespective of how well our treatments mimicked natural microbial communities, our analysis of eight common garden experiments permits assessment of the consistency across time and space of plant genetic effects on their associated microbial communities.

### Host effects on the microbiota

Our experiments provided a unique opportunity to investigate associations between host genetic variation and their resident microbiomes, within the context of environmental variation across time and space. We computed the proportion of variance explained by the host genotype (hereafter heritability or *H^2^*) based on simple unconstrained principal coordinates (PCoA) within each experiment. Within each experiment, we found significant heritability of components of the microbial communities (Extended Data Table 1), suggesting that genetic variation in the host significantly impacts at least a fraction of the microbiota, in line with results of previous studies [8–11,19,20].

Significant heritability of principal components could arise from host genotypes exerting weak control over a large number of community members, or by targeting a few microbes that then influence the relative abundance of others through microbe-microbe interactions. Random-effects linear modeling of log-transformed OTU counts revealed significant genotypic effects (with the 95% confidence interval of heritability not overlapping for between 10.52 and 22.65% of all OTUs, depending on the site and year (Fig. 2A-D and Extended Data Fig. 2A-D). Thus, the influence of the host appears focused on relatively few OTUs. We found no evidence that either fungal or bacterial communities are systematically more impacted by host effects than the other (Fig. 2A-D and Extended Data Fig. 2A-D), nor that mean relative abundance was strongly correlated with OTU heritability (Extended Data Fig. 3).

**Fig. 2.**
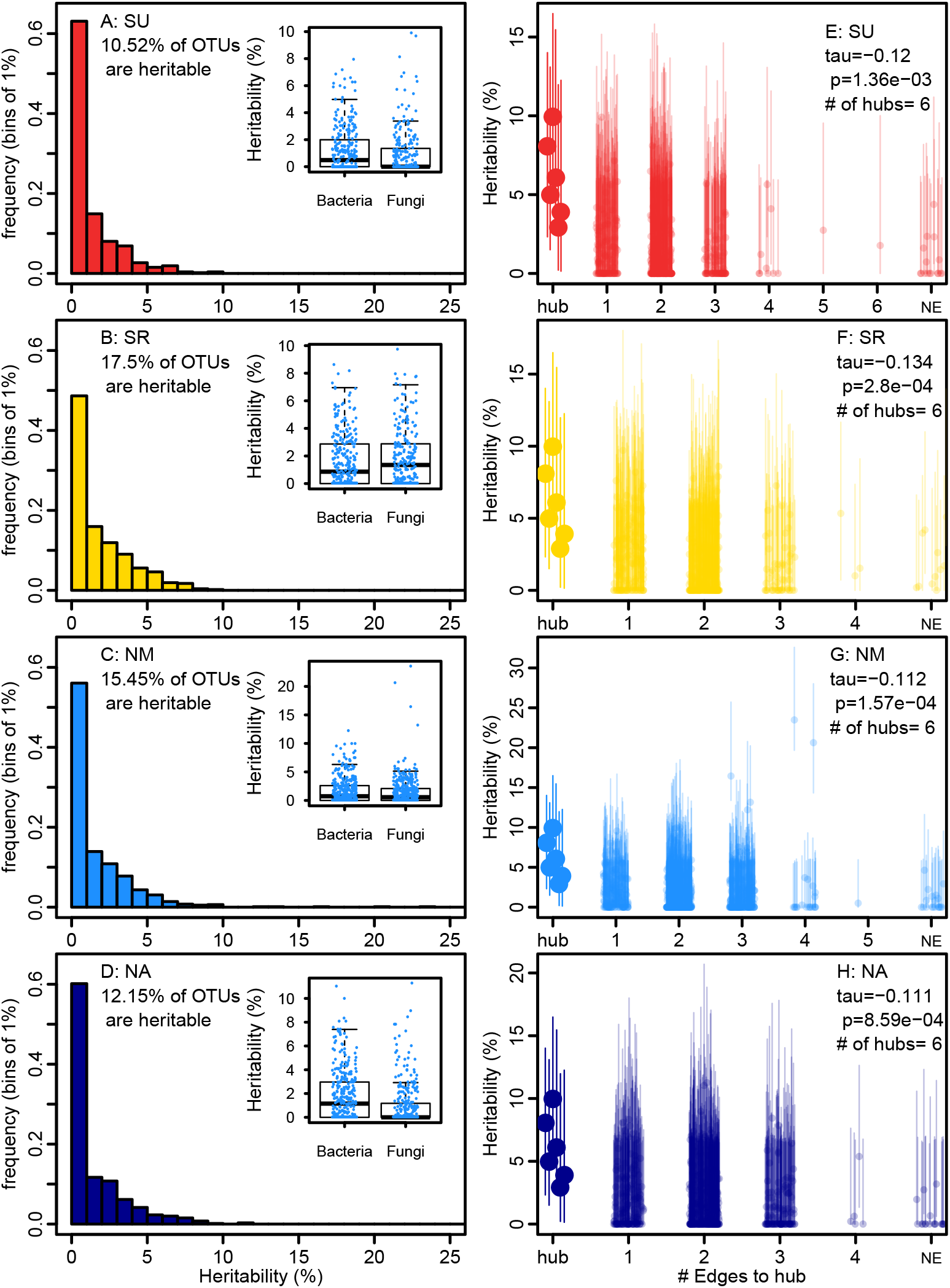
The effect of host genetic variation on the microbial community targets relatively few OTUs and percolates through hubs. This figure corresponds to observations in the set of four experiments sampled in 2013, see Extended Data Figure 3 for experiments performed in 2012. **A-D:** Each frame presents the distribution of heritability estimates for individual OTUs in one site. In each frame, the inset graph is a box and whiskers plot contrasting the heritability (y-axis) of bacterial (B) and fungal (F) OTUs. **E-F:** The heritable hubs are represented by large dots, at a distance of 0 (hub). The other OTUs are represented by smaller dots and the x-axis represents their distance to the nearest heritable hub(s) within the sparse covariance networks. The number of heritable hubs detected in each experiment is indicated in the legend. The correlation coefficients presented are Kendall rank correlations calculated for OTUs with a distance to the heritable hub(s) above 0. NE stands for “no edge”.

Having found that host effects are concentrated on a small proportion of OTUs, we investigated the possibility that these heritable OTUs trigger a broader community level change in the microbiota. First, we computed networks of microbe co-occurrence for each experiment. We explored the ecological importance of heritable OTUs by computing networks of microbe co-occurrence for each experiment using the SPIEC-EASI pipeline [21]. Although our networks included both fungal and bacterial OTUs, most microbe-microbe interactions occurred within each domain, with an average of only 8.71 [min=6.94, max=10.38]% of edges connecting fungal and bacterial OTUs. We quantified the ecological importance of OTUs using two common characteristics of nodes in a network (“Degree” and “Between-ness centrality”) [11], defining ecologically important “hubs” in each network as OTUs in the 95% tail of both of these statistics (Extended data Fig. 4). We identified on average 16.5 microbial hubs per experiment (ranging from 11 to 24), representing 77 unique OTUs across all eight experiments. These hubs were connected to an average of 20.09 [min=14.50, max=25.23]% of the edges in the networks, indicating that they are likely important in structuring the microbial community. In addition, hubs were involved in proportionally more interactions between fungi and bacteria than the rest of the community (Extended Data Table 3).

Next, we asked whether heritable OTUs are more likely to be ecological hubs, because this could open the door to community-level impacts. Across all eight experiments, we detected 23 OTUs that were both heritable and hubs at least once (Extended Data Table 2, Supplementary Table 2). This represents a significant enrichment of hub OTUs amongst heritable OTUs (Wilcoxon rank sum test: N=8, W=57, *p*-value= 0.00699), suggesting that host effects on the microbiota preferentially influence the relative abundance of ecologically important microbes. In fact, hub OTUs were often among the OTUs with the highest heritability within each experiment.

Finally, we sought evidence of community level impacts of heritable hubs by mapping heritability onto the ecological network. In six out of eight experiments, we observed a significant negative relationship between heritability and the distance (number of network edges) to the nearest heritable hub (combined *p*-value=4.104e^−15^, using Fisher’s method for combining *p*-values)[22](Fig. 2E-H and Extended Data Fig. 2E-H). This suggests that host genetic variation most strongly affects a few microbial hubs that then influence other microbes, most likely through microbe-microbe interactions.

Not only do heritable hubs have an impact that appears to percolate through the microbial community, they tend to be widely distributed among accessions, sites and years. We were able to identify 278 fungal and bacterial OTUs that were found in at least 50% of samples in all experiments. Interestingly, OTUs that were heritable hubs at least once were over-represented in this core microbiota (χ^2^=34.68, df=1, *p*-value=3.891e-9). This was not an artifact of their being widespread; significant heritability estimates were detected across the entire range of prevalence, with prevalence of an OTU explaining less than 1.4% of OTU heritability across all experiments (F-statistic=29.48, df=4176, *p*-value=5.964e-08, Extended Data Figure 5). Thus, ecologically important OTUs with greatest associations to host genotypes were unusual in being widespread among plants in multiple experiments. Host effects on the fungal OTU #8 (hereafter F8) are especially important; this OTU was heritable in five out of the seven experiments in which it was a hub (Extended Data Table 2), suggesting that natural variation in *A. thaliana* influences its microbiota with some consistency across environments. The widespread prevalence of these heritable hubs suggests that variation at particular host genes associate with particular hubs across time and space, potentially providing a means to impact the microbiota in a robust fashion.

### Variation in performance of host genotypes explained by their influence on microbial hubs

The extent to which natural variation among host genotypes in their associated microbes translates into fitness differences has yet to be determined. Our experiments included additional replicates of all genotypes that were left to flower and mature in the field. We harvested mature stems in early summer and used high-throughput image analysis to estimate lifetime seed production (LSP) from mature stem size, using an independently validated method (Extended Data Fig. 6) [23]. We observed that plant LSP estimates were positively correlated across experiments (Extended Data Fig. 7), suggesting fitness variation among accessions was relatively consistent across sites. We therefore asked whether host effects on microbial hubs contributed to some genotypes producing more seeds across all environments investigated. Specifically, we used random intercept models to estimate genotype effects on both heritable microbial hubs and LSP in a series of analyses that jointly considered all eight experiments and investigated the relationship between these two effects (see methods “Heritable hubs and LSP across environments”).

We found that the host genotype explained, on average, 6.88% (with a 95% confidence interval [5.52, 8.34]) of our estimate of plant LSP. Host genotype effects (blups) on the relative abundances of 19 of our 23 heritable microbial hubs were similarly modest, explaining up to 4% of the variation (Fig. 3A, four heritable hubs were not detected in more than 2 experiments and were removed for this analysis). In order to estimate genetic correlations between host genotype effects on LSP and on microbial hubs, we performed a multiple regression. After using model selection to identify significant relationships, we detected positive correlations between accession effects on LSP and accession effects on three heritable hubs, F8, B38 and B13, as well as a negative correlation between accession effects on LSP and accession effects on F5 (Fig. 3B). The variation explained by host genotype on the relative abundances of microbial hubs explained 12.4% of the host genotype effects on LSP.

**Fig. 3.**
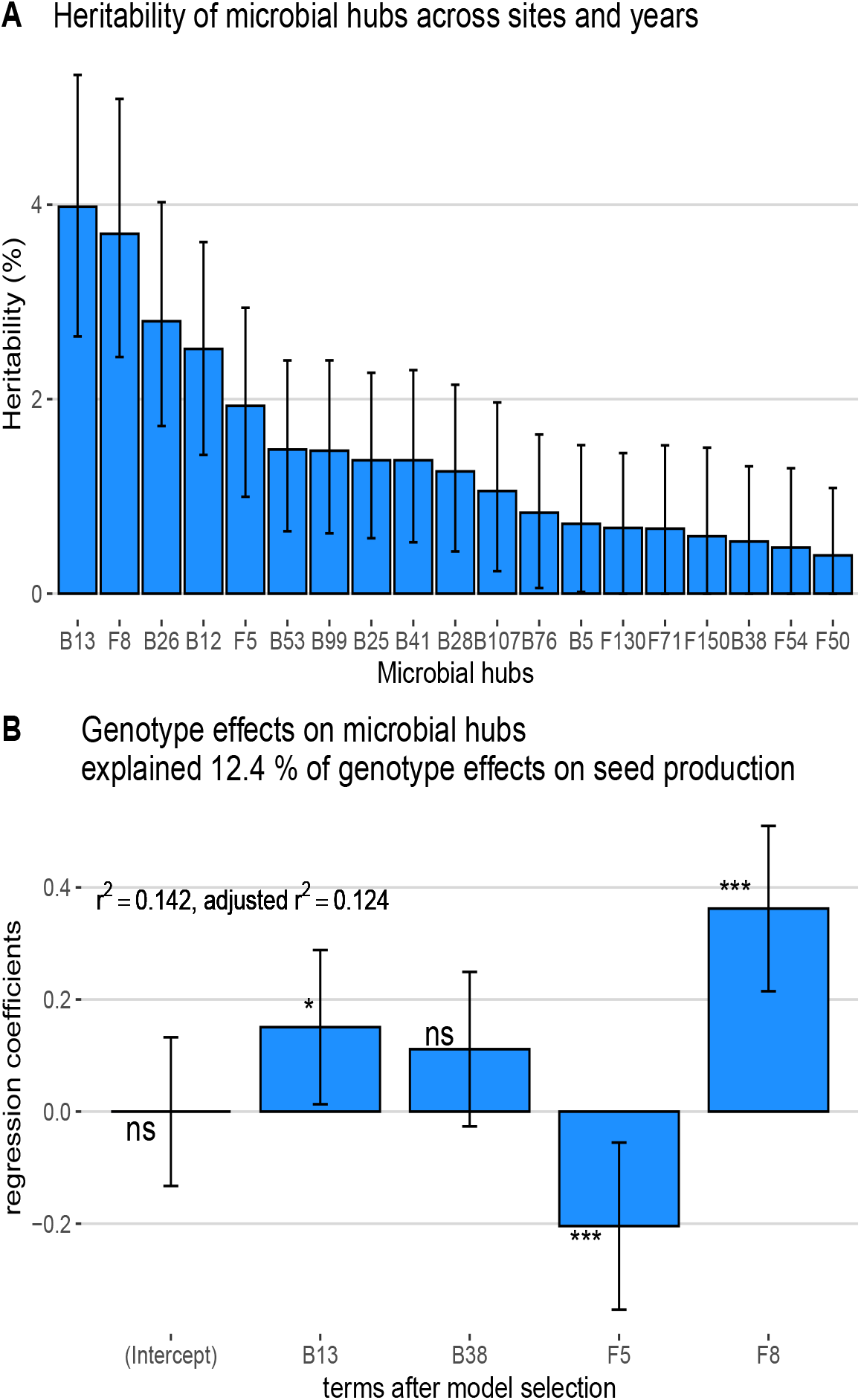
Relationship between host genotype seed production and influence on microbial hubs across sites and years. **A**. Proportion of heritable hub relative counts explained by host effects across all sites and years. **B**. Coefficients for the linear regression explaining lifetime see production variation among accession with accession effects on microbial hubs across experiments (after model selection).

These results reveal that a sizable percentage of genetic variance in LSP is shared with genetic variation associated with the relative abundance of a few broadly distributed microbial hubs, consistent with a causal relationship between genotype and LSP mediated by heritable microbial hubs. Of course, the proportion of shared genetic variation between LSP and heritable microbial hubs is unlikely to be equally important across time and space. In fact, in analyses performed on an experiment-by-experiment basis, we found that relationships between host effects on hubs and LSP were stronger in southern Sweden, where we detected significant relationships in both sites and both years (Extended data Table 4).

Overall, our results highlight the importance for plants to control their leaf microbial community and suggest that breeding plants for their effects on specific members of microbial communities has the potential to significantly increase plant productivity.

### Effect of hubs on growth in controlled condition

In an effort to verify the correlations between host performance and the relative abundance of microbial hubs, we returned to the field to collect wild *A. thaliana* leaves [24], cultured approximately 2400 microbial isolates from within these leaves, and sequenced both the 16S RNA gene and gyrase-B. Among heritable hubs, only B38 was successfully cultured; this isolate derived from Vårhallarna, in southern Sweden, and was identified by a 100% match in 16S sequencing (Extended data Table 5). We subsequently performed shotgun whole genome sequencing of B38 which we identified as *Brevundimonas sp*. The assembled and annotated genome did not identify putative pathogenic or virulence genetic factors present in the genome.

To test the effects of B38 on host growth, we grew Arabidopsis plants of an accession (#6136) from the South of Sweden chosen to have intermediate relative abundance of B38 in the field. Plants were grown under sterile conditions in ½ MS media under long day conditions in the growth chamber, with and without B38 inoculation. Approximately two weeks after germination, over 600 plants were randomly selected for either drip inoculation with the control or B38 inoculum, and measured for surface area growth over the following two weeks. Accounting for variation in plant growth among trials and plates within trials, we found that plants treated with B38 grew 5.375 (standard error=1.973) mm^2^ larger than control plants (*F*=7.3981, df=1, *p*-value=6.7e^−3^) between day 7 and 14, corresponding to a 10.22% growth increase.

The microbial hubs could in principle influence host fitness directly, for example by contributing to growth, or indirectly through their influence on other beneficial members of the microbial community [25]. Here we show that B38 directly improves host growth over early life stages in isolation from the rest of the microbial community. This result is consistent with our field observations, where we found a positive correlation between genetic variation associated with B38 and with LSP, suggesting that in this instance the correlation is causative. The possibility of additional indirect interactions in the field cannot, of course, be excluded.

### Mapping the genetic bases of consistent variation in the relative abundances of microbial hubs across experiments

Our observation that host control of the relative abundances of four microbial hubs explains ~12% of variation in LSP among Arabidopsis genotypes grown in 8 field trials suggests the potential to reveal host genes that can enhance plant performance in the presence of microbes, particularly across environments. Towards this end, we performed genome-wide association mapping for host genotype effects on microbial hubs (N=19) and LSP across all experiments. Despite significant differences among accessions, GWAs yielded few peaks with *p*-values below accepted significance thresholds after correction for multiple testing. Specifically, we found only two significant associations, both for microbial hub B41. The first is located on chromosome 1 at position 29909876 in AT1G79510 annotated as a pseudo-gene. The second is on chromosome 4 on positions 15704377, 15704472 and 15704478. These consecutive SNPs are located between *YUC-1* (AT4G32540), involved in auxin biosynthesis, and *LEUNIG* (AT4G32551), involved in the development of the leaf blade and floral organs.

A potentially more powerful strategy to detect minor QTL involves computing local association scores along the genome. The assumption underlying this method is that neighboring markers in linkage disequilibrium with causal mutations will also carry association signals; thus, aggregating *p*-values increases power [26]. This method identified 340 non-overlapping loci (hereafter QTLs), with sizes ranging from 93 to 150,926 bp including a total of 25,529 SNPs. Out of the 340 QTLs, only 27 included SNPs associated with multiple traits (Supplementary Table 3), suggesting a modest level of pleiotropy.

To investigate functions underlying these associations, we tested pathway and GO term enrichment (Biological processes only)[27,28]. Using a combination of methods accounting for multiple testing, overlapping gene lists, and the potential aggregation of functions and associations along the genome [29–32], we identified 29 enriched GO terms related to biological processes across 16 traits (Supplementary Table 4 and 5), including four genes involved in the response to virus (GO:0009615) and nematodes (GO:0009624), hypersensitive response (GO:0009626) and response to chitin (GO:0010200), all of which are related to interactions with other organisms. Three enriched GO terms directly concern auxins and their transport (GO:0009926, GO:0010540, GO:0009734); auxins have previously been documented to contribute to shaping plant interactions with beneficial bacteria [33,34]. Specialized metabolites also appear involved in shaping the relative abundance of microbial hubs. Indeed hub B107 is associated with genes in the geranylgeranyl diphosphate metabolism (GO:0033385), the universal precursor of monoterpenes, which are volatile compounds with anti-microbial properties, [see 35 for a review] that potentially shape within rosettes microbial communities. In addition, loci associated with B76 are enriched in genes related to specialized metabolite biosynthesis (GO:0044550) and genes involved in the synthesis of sinapoyl glucose and sinapoyl malate (PWY-3301), an intermediate in the synthesis of phenylpropanoids. Genes involved in the synthesis of glucosinolates from phenylalanine (#11 Bz [36] aka glucotropaeolin, PWY-2821) and hexahomomethionine (specifically #69 mSOo [36] aka 8-(methylsulfinyl)octyl-glucosinolate PWYQT-4475) are also enriched in loci associated with B5 and F71, respectively.

The functions highlighted by our analysis are in line with other studies suggesting the involvement of specialized metabolites, auxins and the immune system in influencing the leaf microbial communities [37,38]. Our analysis also highlights less obvious players, like growth lipid metabolism and brassinosteroids (Supplementary Table 5). This is especially true with regard to beneficial members of the community. For example, loci associated with the relative abundance of the beneficial microbial hub B38 are enriched for transition metal ion transport (GO:0000041), response to carbohydrates (GO:0009743), and fatty acid biosynthesis (PWY-4381).

### Plant specialized metabolites correlated with microbial hub abundance

Our biological processes and pathway enrichment analysis suggest that specialized metabolites are involved in shaping microbial hubs. To support this result, we quantified 20 abundant compounds using untargeted metabolomics in a subset of the field samples in which we characterized the rosette microbiome. We found that the relative abundance of 14 out of 19 hubs were significantly correlated with at least one of 11 specialized metabolites (after correction for multiple testing), six of which displayed significant heritability in the field across sites ranging from 1 to 38% (Extended Data Fig. 8A & B).

The molecule known as #69 mSOo (here 260_GSL_8MSO) displayed the strongest relationship with multiple microbial hubs in the field (Extended Data Fig. 8A, Extended Data Table 6), as well as significant heritability under field conditions (Extended Data Fig. 8B). However, the variation among accessions of this abundant glucosinolate was less evident in the greenhouse and in sterile conditions (Extended Data Fig. 8B), leaving open the possibility that the correlation is induced by one or more of the microbial hubs. In contrast, other molecules significantly related with the abundance of microbial hubs in the field across experiments (354_C_Cy-GRGF_785 and 358_F_R-K-R_577, Extended Data Table 6) are heritable in all conditions, and variation among accessions in the field is positively correlated with the variation among accessions in the greenhouse. This suggests that these flavonoids are constitutively and consistently produced by accessions and influence microbial hubs in a manner that is robust to heterogeneity among field experiments.

## Conclusion

In this study, we show that not only does host genetic variation influence the microbiome, but it does so in consistent ways. Host genotype effects are centered on ecologically important hub species, and percolate through the microbial community, most likely as a result of microbe-microbe interactions. Our replicate field experiments were likely instrumental in allowing us to reveal consistent host effects on the leaf microbiome via common and widespread hub species.

Furthermore, we found that the influence of host genetics on a handful of prevalent microbial hubs has a far-reaching impact on the community, associated with a substantial fraction of the variation in our fitness estimates among accessions. Although these relationships are correlational, we were able to culture one of the identified hubs and confirm a direct positive effect on host fitness experimentally.

Understanding how host performance or fitness components are influenced by their ability to shape microbial communities could provide a basis for breeding crops favoring microbes that are beneficial both to growth and resistance to pathogens. We successfully mapped variation in host microbe interactions using genome-wide association, and our results suggest that natural and artificial selection can act on plant traits such as leaf specialized metabolites, auxins and the immune system to improve plant performances through effects on microbial communities [39,40]. In addition, we found that at least some plant metabolites are expressed in a consistent manner that is robust to variation among our experiments and correlates with the relative abundance of microbial hubs. Our results therefore suggest that ongoing efforts to harness the microbiome for agricultural purposes can be successful and highlight the value of explicitly considering abiotic variation in those efforts.

## Methods

### Field experiments

This study uses a set of 200 diverse accessions (inbred lines, Supplementary Table 1) that were previously re-sequenced [14]. The seeds were produced simultaneously in the greenhouse of the University of Chicago under long day conditions, except for a 12-week vernalization period at 4°C, required to induce flowering. The seeds for the common garden experiments were cold stratified in water at 4°C for 3 days before being planted in trays of 66 open-bottom wells, each measuring 4 cm in diameter. The soil used was a 90:10 mix of standard greenhouse soil and soil from each of the four sites in which the experiments were installed:

– SU: Ullstorp (Agricultural field, lat: 56.067, long: 13.945)
– SR: Ratchkegården (Agricultural field, lat: 55.906, long: 14.260)
– NM: Ramsta (Agricultural field, lat: 62.85, long: 18.193)
– NA: Ådal (South facing slope, lat: 62.862, long 18.331)

Each experiment included 3 complete randomized blocks including 8 replicates per accession. Experiments were sown in pairs (2 in the North and 2 in the South) over 6 days, corresponding to the sowing of one block a day, alternating between the 2 experiments (between August 7th and 12th in the North, and between August 31st and September 5th in the South). The trays were placed in a common garden the morning after sowing under row tunnels to avoid disturbance by precipitation and to favor germination (on the campus of Mid Sweden University and Lund University, in the North and in the South, respectively). Trays were watered as needed and missing seedlings were transplanted between cells within blocks and then thinned to one per cell after 9 days. Seventeen days after sowing, trays were laid in the field in their final location over tilled soil. For each experiment, the blocks were laid across the most obvious environmental gradient (exposition, shading, slope, soil humidity…). The pierced bottom of the cells allowed the roots to grow through and reach the soil, as was verified upon harvest. The same protocol was followed in 2011 and 2012.

### Sample collection and processing

The rosettes used to characterize the microbial community were harvested in the spring of 2012 and 2013 only a few days after the plants were exposed, following snow melt. We harvested 2 randomly selected replicates per accession in each experimental block. Upon harvest, the roots were removed and the rosettes were washed twice in successive baths of TE and 70% ethanol to remove loosely attached microbes from the leaf surface. The rosettes were then placed in sealed paper envelopes and placed on dry ice. The rosettes were kept at − 80°C until lyophilized. Freeze-dried rosettes were then transferred to 2 ml tubes along with 3 2mm silica beads. For 2 successive years, the tubes were randomized and separated in 34 and 46 sets of 96 tubes, respectively. Our randomization strategy maintained approximately the same number of tubes from each of the 12 experimental units (3 blocks in 4 experiments) in order to avoid confounding biologically meaningful effects. We powdered the samples using a Geno/Grinder® (from Spex SamplePrep, USA, NJ) for 1min at 1750rpm, before transferring 10 - 20 mg to 2ml 96-well plates, along with two zirconia/silica beads (diameter = 2.3mm), for DNA extraction.

### DNA extraction

DNA extraction started with 2 enzymatic digestions to maximize yield from Gram-negative bacteria [41]. First, we added 250μl of TES with 50 units.μl^−1^ of Lysozyme (Ready-Lys Lysozyme, Epicenter) to each well. The plates were then shaken using the Geno-Grinder for 2 min at 1750 rpm, briefly spun and incubated 30 min at room temperature. Second, we added 250μl of TES with 2% SDS and 1 mg.mL^−1^ of proteinase K. The plates were then briefly vortexed and incubated at 55°C for 4 hours. The protocol then followed [42], adapted to the 96-well plate format and automated pipetting on a Tecan Freedom Evo Liquid Handler. We added 500 μl of Chloroform:Isoamyl Alcohol (24:1), pipette mixed, and centrifuged the plates at 6600 g for 15 min. We transferred 450 μl of the aqueous supernatant to a new plate containing 500μl of 100% isopropanol. The plates were then sealed, inverted 50 times, incubated at −20°C for 1 hour, and centrifuged at 6600 g for 15 min. The Isopropanol was then removed and the pellets were washed twice with 500 μl of 70% Ethanol, dried and re-suspended in 100 μl of TE. After 5 min incubation on ice, the plates were centrifuged 12 min at 6600 g and the supernatant was pipetted into a new plate.

### PCR and Sequencing

To describe the microbial communities, we amplified and sequenced fragments of the taxonomically informative genes *16S* and *ITS* for bacteria and fungi, respectively. For bacteria we amplified the hypervariable regions V5, V6 and V7 of the *16S* gene using the primers 799F (5′-AACMGGATTAGATACCCKG-3′) and 1193R (5′-ACGTCATCCCCACCTTCC-3′) [9,43]. For fungi, we amplified the ITS-1 region using the primers ITS1F (5′-CTTGGTCATTTAGAGGAAGTAA-3′) [15] and ITS2 (5′-GCTGCGTTCTTCATCGATGC-3′) [44]. To the 5′ end of these primers we added a 2bp linker, a 10bp pad region, a 6bp barcode and the adapter to the Illumina flowcell, following [45]. The appropriate linkers were chosen using the PrimerProspector program [46]. The PCR reactions were realized in 25 μl including: 10 μl of Hot Start Master Mix 2.5x (5prime), 1μl of a 1/10 dilution of the DNA template, 4μl of SBT-PAR buffer, and 5 μl of the forward and reverse primers (1 μM). The SBT-PAR buffer is a modified version of the TBT-PAR PCR buffer described in [47] with the trehalose replaced by sucrose (Sucrose, BSA, Tween20). The PCR program consisted of an initial denaturing step at 94°C for 2’30”, followed by 35 cycles of a denaturing step (94°C for 30”), an annealing step (54.3°C for 40”), and an extension step (68°C for 40”). A final extension step at 68°C was performed for 7’ before storing the samples at 4°C. For each plate, the PCRs were performed in triplicates, pooled, and purified using 90 μl of a magnetic bead solution prepared and used following [48]. The purified PCR products were quantified with Picogreen following the manufacturer’s instruction [49] and pooled into an equimolar mix. Between 5 and 7 plates (480 to 672 samples) were pooled in each MiSeq run. If the bioanalyzer traces for pooled libraries showed only one dominant peak, they were sequenced directly following the standard MiSeq library preparation protocols for amplicons. In cases where the bioanalyzer trace presented peaks for smaller fragments (remaining primers, primer dimers, small PCR products), the libraries were first concentrated 20X on a speedvac (55°C for 2 to 3 hours), purified with 0.9 volume of magnetic bead solution, and/or size selected using a Blue Pippin (range mode between 300 and 800 bp).

The sequencing was performed using MiSeq 500 cycle V2 kits (251 cycles per read and 6 cycles of index reads twice), using a loading concentration of 12.5pM for *ITS* fragments and 8pM for *16S* fragments following the standard Illumina protocol. Sequencing primers were designed and spiked in following [45]. The sequencing primer for the first read of *16S* fragments was prolonged into the conserved beginning of the fragment amplified to reach a sufficient melting temperature. This primer modification produced no change in the Blast results of the primers against the GreenGene database. A total of 11 sequencing runs were performed for each of the fungal and bacterial communities.

### Sequence processing and clustering

The demultiplexed fastq files generated by MiSeq reporter for the first read of each run were quality filtered and truncated to remove potential primer sequences and low quality basecalls using the program cutadapt [50]. The reads were then further filtered and converted to fasta files using the FASTX-Toolkit (-q 30 −p 90 −Q33). The fasta files for each run were then de-replicated using AWK code provided in the swarm git repository (https://github.com/torognes/swarm)[16]. The resulting de-replicated fasta files were filtered for PCR chimeras using the vsearch uchime_denovo command (https://github.com/torognes/vsearch). The de-replicated fasta files for each run were then combined and further de-replicated at the study level. The fasta files were then used as input for OTU clustering using swarm (−t 4 −c 20000). The clustering identified 150,412 and 251,065 OTUs for the fungal and bacterial communities, respectively. The output files were combined into two separate community matrices using a custom python script (available at https://bitbucket.org/bbrachi/microbiota.git). The taxonomy of each OTU was determined using the quiime2 2019.1 v8 feature classifier trained on the UNITE V6 and SILVA 119 database for Bacteria and Fungi, respectively [51,52].

### Count table filtering

The count tables obtained for both the bacterial and fungal communities were filtered in successive steps by removing:

1. samples corresponding to empty wells and additional plant genotypes present in the experiments sampled by mistake (leaving 7476 and 7240 samples for the fungal and bacterial count tables, respectively).
2. samples with less than 1000 reads (leaving 6678 and 6819 samples for the fungal and bacterial count tables, respectively)
3. OTUs represented by less than 10 read in 5 samples (leaving 1381 and 993 OTUs for the fungal and bacterial count tables, respectively)
4. for the bacterial community, OTUs assigned to plant mitochondria (leaving 993 OTUs in the bacterial count table, no OTUs assigned to plant mitochondria)
5. for a second time, samples with less than 1000 reads (leaving 6656 and 6783 samples for the fungal and bacterial count tables, respectively).

The final count tables used in the study included 993 OTUs and 6783 samples for the bacterial communities and 1381 OTUs and 6656 samples for the fungal community.

### Differentiation of the microbial communities among sites and years

This analysis was performed for the fungal and bacterial communities independently, including all samples and only OTUs with read counts above 0.01% of total read counts (after the filtering described above) across sites and years. To investigate how the microbial communities differed among sites and years, we performed a constrained ordination on log transformed read counts using the capscale function in the R-package Vegan [53] and following [54]. The log transformation offers the advantage of removing large differences in scale among variables. The capscale function performs canonical analysis of principal coordinates, an analysis similar to redundancy analysis (rda), but based on the decomposition of a Bray-Curtis dissimilarity matrix among samples (instead of euclidean distance in the case of rda). This allows identification of the dimension that maximized the variance explained by components, while discriminating groups of samples, here sites and years [54].

### Core microbiota

In order to define a core microbiota, we counted, for each OTU, the number of site/year combinations in which it was prevalent. We defined “prevalent” as being present in at least 50% of the samples in a given site/year. We performed this analysis using count tables for each experiment with the filtering described in the previous paragraph. Therefore, for an OTU to be designated as a member of the core microbiota, it needed to have non-zero counts in more than 50% of the samples within each site/year combinations and, due to previously described filtering, needed to be represented by at least 10 reads in 5 of those samples across all site/year combinations (see “Count table filtering”).

### Heritability of the microbiota

In this analysis, count tables were split per site and year before filtering for OTUs represented by more than 0.01% of the reads (after the filtering described in the section “Count table filtering”) for each of the bacterial and fungal communities. The resulting 16 count tables were normalized to 1000 reads per sample and used to calculate 16 Bray-Curtis pairwise dissimilarity matrices among samples. These matrices were then decomposed into 10 principal coordinates. For each component we estimated broad sense heritability (hereafter H2), *i.e.* the proportion of variance explained by a random intercept effect capturing the identity of the accessions present in the experiment (plate effects had limited impact on H2 estimates but were included in the models). Mixed models were fitted using the function lmer in the lme4 R package [55]. We computed 95% confidence intervals using 1000 bootstraps, and components were considered to have significant H2 when their confidence intervals did not overlap 0 (lower bound of the confidence interval ≥ 0.01).

### Heritability of individual OTUs

This analysis was also performed per site, year and community, as in the microbiota H^2^ estimation analysis. In this analysis, counts were transformed to centered log-ratios using a dedicated function in the R package mixOmics [56,57]. H^2^ estimates and confidence intervals were computed for individual OTUs using the method described in the previous paragraph (without the plate effect). H^2^ estimates for our estimate of LSP (see below) were estimated the same way using a box-cox transformation.

### Microbe–microbe interaction networks

Microbe-microbe interaction networks were computed for the fungal and bacterial communities together, using the count tables per site/ year and filtering OTUs represented by less than 0.01% of the reads within each community. The count tables were then combined into the same table and analyzed using the SPIEC-EASI (v1.1) pipeline [21]. This method computes sparse microbial ecological networks in a fashion robust to compositional bias and uses conditional independence to identify true ecological interactions, meaning that a connection between 2 OTUs will be significant when one provides information about the other, given the state of all other OTUs in the network. This means that covariance among OTUs induced by micro-environmental and host genetic variation is controlled. SPIEC-EASI was run using the neighborhood selection framework and model selection was regularized with parameters set to a minimum lambda ratio of 1e^−2^ and a sequence of 50 lambda values (see documentation for SPIEC-EASI and the huge R package, which provides regularization functions)[58].

### Network statistics

The inferences of microbe-microbe ecological interactions inferred using SPIEC-EASI were passed to the igraph package [59], which was used for enforcing simplicity of graphs (no loops or duplicated edges), computing degree and betweenness centrality of vertices, computing distances between vertices, and plotting. Within each of the 8 networks thus computed, hubs were defined as OTUs with degree and betweenness centrality both in the 5% tail of their respective distributions. We then checked the overlap between heritable OTUs and hubs, and the over-representation of heritable OTUs among hubs was tested using a simple χ^2^ test across all site/year combinations. The relationship between distances to heritable hubs (OTUs that are both hubs and have significant H^2^) and heritability was investigated using Spearman’s rank correlation coefficient. Distances were calculated as the number of edges between OTUs and the closest heritable hub in the network. OTUs not connected to heritable hubs were assigned a distance equal to one more than the maximum distance observed for OTUs connected to heritable hubs.

### Estimation of seed production

The experiments each included 8 replicates per block per accession (24 replicates per experiment). While we harvested 2 replicates per block (6 replicates per experiment) for microbiota analysis, the remaining plants were left to grow, flower and produce seeds in the field. We harvested the mature stems of all remaining plants at the end of the spring, when all plants had finished flowering and siliques were mature, and stored them flat in individual paper envelopes. We estimated lifetime see production (LSP) by the size of the mature stems. After removing remaining traces of roots and rosettes, each mature plant was photographed on a black background, using a DSLR camera (Nikon 60D) mounted on a copy-stand and equipped with a 60mm macro lens (Nikon 60mm). The photographs were segmented (using custom scripts in R based on the EBimage package [60] to isolate plants from the image background and estimate the total surface of the image they occupied.

We validated this method with mature plants harvested from a previous experiment that was planted in NM in fall 2010, and that included the 200 accessions used in this study. We counted siliques and estimated the average silique size for 1607 mature stems that were also photographed. The total silique length produced per plant (number * average size) was highly correlated with our size estimates based on image analysis (Spearman’s rho=0.84) and displayed a clear linear relationship.

### Relationship between host effects on microbial hubs and fecundity

To investigate the relationship between host genotype effects on heritable hubs and LSP in each experiment, we computed estimates of accession effects (Best unbiased linear predictors or BLUPs) for both log-ratio transformed heritable hubs and box-cox transformed LSP estimates. We then fitted multiple regressions for each site/year combination aiming to explain LSP variation among accessions with their influence over microbial hubs and following **eq. 1**.

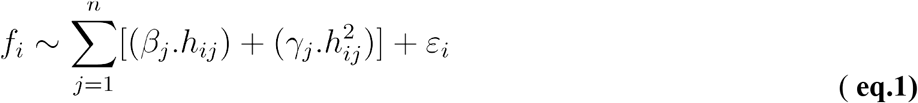

where *f_i_* is the LSP estimate of the i^th^ accession (blup), *h_ij_* is the effect of the i^th^ accession on the j^th^ hub. *β_j_* is the regression coefficient for the j^th^ hub (*h_j_*) and *γ_j_* is the regression coefficient for the j^th^ hub squared. 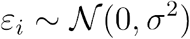 captures residual variance per accession. We then performed forward/backward model selection to obtain the final models presented in (Extended Data Table 4).

### Heritable hubs and LSP across environments

We next investigated host effects on heritable hubs and LSP across all 8 experiments. Similarly to previous analyses, count tables were split per site and year before filtering for OTUs represented by more than 0.01% of the reads (after the filtering described in the section “Count table filtering”) for each of the bacterial and fungal communities. The resulting 16 count tables were then combined into one before fitting a mixed-model following eq. 2:

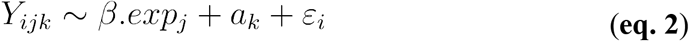

where *Y_ijk_* are the transformed counts for a heritable hub measured the i^th^ time in experiment 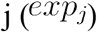 (N=8, four sites and two years) and for accession k (N=200), *β* is the vector of fixed experiment effects (N=8) and 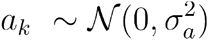 is a random intercept estimated by restricted maximum likelihood for each accession.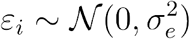 captures the residual variance.

Microbial hub heritability (H^2^) across experiments was estimated as the percentage of variance explained by the random accession intercept:

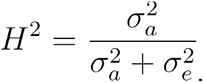

LSP data was analyzed the same way, except we performed Box-Cox transformation of the data. The lambda parameter for the Box-Cox transformation was estimated using the same model, but without the random accession term.

For both heritable microbial hubs and LSP, we retrieved random intercept accession effects (BLUPS) and fitted a multiple linear regression following:

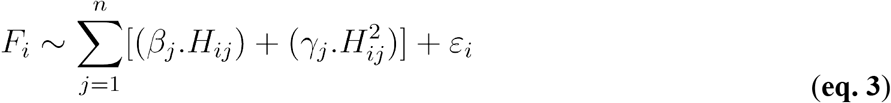

where *F_i_* is the effect of the i^th^ accession (N=200) on LSP (across all experiments), *H_ij_* is the effect of accession i on hub j across all experiments, 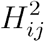 is the squared effect of accession i on hub j. *β_j_* and *γ_j_* are the corresponding regression coefficient for hub j and 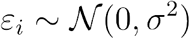 captures the residual variance per accession. The final model was obtained after backward/forward model selection based on AIC.

### Isolation, culture and identification of microbial hubs

#### Bacteria sampling from wild A. thaliana plants

We collected 2 leaves from 10 plants at 5 locations in Sweden (Extended data Table 5). The leaves were first cleaned by rinsing individually in ddH2O, and subsequently surface-sterilized by dipping 70% EtOH for 3-5 seconds. The leaves were ground in individual 1.5 mL tubes. The leaf material was stored in 20% glycerol at −20°C. Wild *A. thaliana* microbial isolates were collected using modified methods that were previously described (Bai et al 2015). Briefly, the leaf and glycerol mixture was plated on nine distinct media, including; R2A, Minimal media containing Methanol, Tryptic Soy Agar, Tryptone Yeast extract Glucose Agar, Yeast Extract Manitol Agar [24]; 0.1 Tryptic Soy Agar [61]; Potato Dextrose Agar, 0.2 Potato Dextrose Agar, and Malt Extract Agar [62,63]. Colonies were picked over the next 14 days, restreaked, and grown in liquid media in an orbital shaker for 1-4 days. A portion of the inoculum was saved in 15-20% glycerol, and the rest of the liquid culture was pelleted by centrifugation and decanted for DNA extraction. We performed a double enzymatic digest for all isolates, which was performed using the Tecan: 30 minute incubation with 350 U Ready-Lyse Lysozyme and 245 U RNase A (QIAGEN, Germantown, MD) in 250ul TES (10 mM Tris-HCl pH ~8, 1 mM EDTA, 100 mM NaCl), followed by the addition of 2 mg/mL Proteinase K in 250ul TES + 2% SDS and a 4-6 hour incubation at 55C. The SDS-protein complexes were precipitated with.3 volume 5M NaCl and pelleted by a brief centrifugation. The clear supernatant was pipetted into a clean plate, and a standard.5 volume SPRI bead DNA extraction was performed with 2x 70% EtOH washes. Clean DNA was resuspended into MilliQ water. The samples were then amplified for 16S sequencing using the same primers binding regions as previously, 799F and 1193R, and sequenced by either Sanger or Illumina MiSeq (PE 300). Illumina adapters were designed and generated as described by Illumina with internal barcodes to increase sample count capacity per lane [64]. Isolate B38 was identified by 100% match to the B38 representative sequence from the previous analysis.

### B38 whole genome assembly

We used a low-input method for Illumina library prep [Baym]. Briefly, ~2 ng extracted DNA was used in a reduced volume (5ul) tagmentation reaction with TDE1 (incubate 55C for 10 mins, room temperature for 5 mins). The tagmentation reaction was added to a 15 ul PCR reaction, adding the Illumina adapters (Kapa HiFi Hotstart PCR kit KK502, standard Illumina adapters and cycling). The library was cleaned with.8x volume SPRI beads, quantified on the Bioanalyzer, and run on the MIseq2500 using paired end 300 chemistry. Reads were trimmed for adapters (BBDuk, ktrim=r k=23 mink=11 hdist=1 tbo) and quality across a sliding window (k=4, trimq=20). Reads were assembled using SPAdes (-isolate) and annotated with PROKKA. **Plant growths assays with B38**

#### Plant growth

*Arabidopsis thaliana* accession 6136 from Southern Sweden was used in the growth assays. In our field experiments it displayed average relative counts for B38 (rank 102 of 199). The plant assay used slightly modified methods as previously described [65]. The seeds were exposed to chlorine gas for sterilization: in a bell jar with dessicant, an open 1.5 mL tube with seeds was placed next to a 50 mL beaker with 40 mL Chlorox bleach and 1 mL hydrochloric acid, sealed with parafilm, and incubated for 4 hours. Sterilized seeds were subsequently sown on 24-well tissue plates containing 1.5mL of ½ MS media (Murashige & Skoog medium incl. Nitsch vitamins, bioWORLD, Dublin, Ohio) containing 500mg/L MES, pH 5.7 - 5.8. Plates were wrapped in parafilm, and vernalized in the dark at 4°C for 4 days. The plates were individually wrapped with micropore tape to prevent environmental contamination and transferred to a growth chamber with 16 hours of light at 16°C. The plants were treated with either B38 or control inoculum between days 13-15 post-vernalization. The plates were returned to the chamber to grow for another 14 days.

#### B38 inoculation

The B38 isolate grew in R2A liquid media in an orbital shaker until OD_600_=0.2, approximately 3 days. To ensure no environmental contamination, a portion of the inoculum was saved for DNA extraction and subsequent 16S Sanger sequencing verification. The liquid cultures were pelleted by centrifuging 1800 RCF 18C for 7 minutes, decanted and resuspended in 0.1 M MgSO_4_. The plants in each 24-well plate were randomly selected to receive the infection (B38 + 0.1 M MgSO_4_) or control (0.1 M MgSO_4_) treatment. Each plant was drip inoculated using pipettes with 180ul of the selected treatment. The plates were re-wrapped in micropore tape and returned to the growth chamber.

#### Measuring plant growth

We performed 3 trials of 11, 28, and 23 plates, totalling 62 24-well plates. Plants were not treated and removed from the experiment if they had less than 3 true leaves, cracked agar, or failed to germinate, resulting in a total of 1094 plants. The plants were individually photographed immediately before inoculation, then again at 7 and 14 days post-inoculation. The images were processed using a custom script employing cv2 in Python [66], which quantified plant surface area in each well by scaling based on the wells’ size, converting images into binary images, and measuring non-white pixels within each well (i.e. plant surface area). The output images were manually inspected, and any images which failed to be accurately processed were manually measured using the same pipeline described above, but using Image J.

Due to the high humidity of the plates and the drip inoculation, 422 plants showed signs of water log stress. Plants were scored for symptoms of stress induced by water logging (blindly with regard to B38 inoculation) as categorized by translucent/white leaves or stunted growth, and were removed from the experiment. We used a linear mixed model (eq. 4) accounting for variation in plant growth among trials and plates within trials to estimate the effect of B38 inoculation.

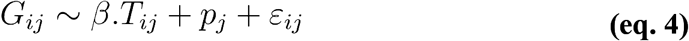

In equation 4, *G_ij_* is the growth of i^th^ plant in the j^th^ plate/assay combination. *β* is the estimate of the treatment effect compared to the controls (intercept) and *T_ij_*, is the treatment (inoculation with a B38 or control solution). 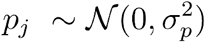 the random intercept effect capturing variation among plates in assays (N=62 plates across three trials).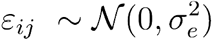 captures the residual variances.

## Genome-wide association mapping

### Single polymorphism calling and filtering

Single nucleotide polymorphisms (SNP) used in this study were generated from the sequences generated in the context of the 1001genome project [67] and published in Long, Q. *et al.* [14]. As pipelines evolve, we re-ran SNP calling to ensure optimal quality. For each sequenced individual, we performed 3’ adapter removal (either TruSeq or Nextera), quality trimming (quality 15 and 10 for 5’ and 3’-ends, respectively) and N-end trimming with cutadapt (v1.9) [50]. After processing, we only kept reads of approximately half the length of the original read-length. We mapped all paired-end (PE) reads to the *A. thaliana* TAIR10 reference genome with BWA-MEM (v0.7.8) [68,69]. We used Samtools (v0.1.18) to convert file formats [70] and Sambamba (v0.6.3) to sort and index bam files [71]. We removed duplicated reads with Markduplicates from Picard (v1.101) (http://broadinstitute.github.io/picard/) and performed local realignment around indels with GATK/RealignerTargetCreator and GATK/IndelRealigner functions from GATK (v3.5) [72,73] by providing known indels from The 1001 Genomes Consortium <u>(1001 Genomes Consortium 2016)</u>. Similarly, we conducted base quality recalibration with the functions GATK/BaseRecalibrator and GATK/PrintReads by providing known indels and SNPs from The 1001 Genomes Consortium.

For variant calling, we employed GATK/HaplotypeCaller on each sample in ‘GVCF mode’, followed by joint genotyping of a single cohort of 220 individuals with GATK/GenotypeGVCFs. To filter SNP variants, we followed the protocol of variant quality score recalibration (VQSR) from GATK. First, we created a set of 191,968 training variants from the intersection between the 250k SNP array [74] used to genotype the RegMap panel and the SNPs from The 1001 Genomes Consortium. Second, this training set was further filtered by the behavior in the population of several annotation profiles (DP < 10686, InbreedingCoeff > −0.1, SOR < 2, FS < 10, MQ > 45, QD > 20) to leave 175,224 training high-quality variants. Third, we executed GATK/VariantRecalibrator with the latter as the training set, an *a priori* probability of 15, the maximum number of Gaussian distributions set at 4, and annotations MQ, MQRankSum, ReadPosRankSum, FS, SOR, DP, QD and InbreedingCoeff enabled. Finally, we applied a sensitivity threshold of 99.5 with GATK/ApplyRecalibration and restricted our set to bi-allelic SNPs with GATK/SelectVariants for a total of 2,303,415 SNPs in the population.

Preparation for use in genome-wide association analysis involved further filtering of individuals and SNPs using Plink1.9 [76,77]. Individuals not included in this study were removed and SNPs with over 5% missing data and with minor allele frequencies below 5% in our collection of accessions were removed.

### Phenotype preparation and association analysis

Association mapping analyses were performed for the 11 heritable microbial hubs for which we estimated host genotype effects across experiment and accession LSP estimates. Association analyses were performed using a classical one trait mixed model accounting for genetic relatedness among accessions (kinship) [78].

In order to take advantage of linkage disequilibrium and gain power by grouping association statistics in contiguous markers, we computed local association scores [26]. We followed the instructions provided by the authors and defined the parameter *X_i_* as the 0.999 quantile of the distribution of*−log(p - value) − 1* rounded to the closest integer for each trait investigated (19 microbial hubs and LSP). The approach highlights regions, which we call QTLs.

The null association model (without fixed SNP effect) from Gemma allows us to estimate SNP-based heritability or pseudo-heritability [79], which is the proportion of variance explained by the random accession effect, accounting for the genetic similarity among accessions. To investigate if the regions highlighted by the local score approach included true positives, we computed SNP based heritability for each trait, each time using three sets of SNPs to compute the kinship matrix: 1) All the SNPs in the genome over 10% frequency, 2) all the SNPs within QTLs identified by the local score approach, and 3) all SNPs not included in the QTLs identified by the local score approach.

### Pathway enrichment analysis

To investigate biological functions associated with LSP of accessions or their influence over microbial hubs, we searched for enrichment in annotated pathways (BIOCYC) and GO categories (Biological processes only) in *Arabidopsis thaliana*. Gene-set enrichment methods are designed for assays that directly assign *p*-values or effects to individual genes (i.e. RNAseq experiments). Here, for each trait, each gene was attributed the largest absolute SNP effect within a distance of 5kb on each side and followed the setRank procedure that accounts for overlapping categories and multiple testing. We set the parameter “setPCutoff” to 0.01 and the “fdrCutoff” to 0.05 [29]. To account for specificities of gene set enrichment in the context of association mapping, we also tested the enrichment of the gene groups identified by setRank using a weighted Kolmogorov-Smirnov score [30] and a permutation scheme accounting for the non-independence of marker effects due to linkage disequilibrium along the genome, as well as the potential clustering of genes with similar function [31,32]. Briefly, enrichment was calculated using a weighted Kolmogorov sum using gene effect rank (and not a gene effect significance threshold)[30]. Enrichments were then tested against an empirical distribution generated from 1e5 permutations. For each permutation, chromosomes are randomly re-ordered and re-oriented and the whole genome is shifted (or “rotated”) by a random number, before re-assigning SNP effects to genes and calculating enrichment for the groups of genes of interest. We considered only categories with an empirical *p*-values below 0.05.

## Untargeted metabolomics

### Plant material and sample preparation

This analysis uses three sets of samples. The first are samples collected from the experiments in Sweden and correspond to a subset of those used for the microbial community. In particular we chose samples from the four experiments established in 2012 and focused on a subset of 50 accessions selected to span the genetic variation among hosts in our mapping population. The second set of samples correspond to 6 replicates of the same 50 genotypes grown in the University of Chicago greenhouse during the summer 2014 under long day conditions (16-hour light period), in standard culture soil. After 28 days, plants were vernalized for three weeks at 4°C and leaf samples were collected after vernalization, immediately flash frozen in liquid nitrogen, freeze-dried and stored at room temperature. The third set corresponds to 3 replicates of the same 50 genotypes, grown on sterile agar medium (Murashige and Skoog with Nitsch vitamins) in individual well plates in a growth chamber with a 16-hour light period (long day condition). Seeds were sterilized by a 70% ethanol bath for 10 minutes, and manipulated under a sterile hood. Samples were collected after 28 days of growth, flash frozen, freeze-dried, and stored at room temperature.

Dried samples from the 3 sets were coarsely ground, and distributed in 18 96-well plates with two ceramic grinding beads per well (10mg per well +/- 2mg). Samples were randomized across all plates to limit confounding of biological effects. In addition, each plate included 16 random samples (1/6) from each experimental unit (greenhouse, sterile, and the 4 field experiments).

### Specialized metabolite extraction and LC-MS analysis

The extraction protocol was designed to extract polar compounds such as glucosinolates and flavonoids. Samples in plates were ground using a Geno/Grinder (SPEX SamplePrep 2010) at 1750 rpm for two minutes. The extraction buffer (70% methanol, 30% water, internal standard: quercetin, 0.0708 mM) was added using a Tecan pipetting robot (100 μl per milligram of dry material). Samples were shaken at room temperature for two hours and filtered on 96-well filter plates (0.45μm) on a vacuum manifold. The flow-through was collected in 96-well plates and stored at 4°C.

Samples were auto injected through a Zorbax SB-C18 2.1 × 150 mm, 3.5 μm column on an Agilent Q-TOF LC–MS with dual ESI (Agilent 6520) with the following parameters: 325 °C gas temperature, 6 L min–1 drying gas, 35 eV fixed collision energy, 35 psig nebulizer, 68 V skimmer voltage, 750 V OCT 1 RF Vpp, 170 V fragmentor, and 3500 V capillary voltage. Mass accuracy was within 2–5 ppm. Samples were eluted with 0.1% formic acid in water (A) and 100% acetonitrile (B) using the following separation gradient: 95% A injection followed by a gradient to 90% A at 1 min, 45% A at 6 min, 100%B at 6.5 min with 4 min hold and 3 min equilibration. An external standard (sinigrin, 1mM) was run 4 times before each plate and one time every 20 samples to monitor and maintain run quality. Compounds were characterized using retention times and fragmentation patterns of chromatograms with automatic agile integration in Agilent Mass Hunter Software (Qualitative Analysis B6 2012) and fragments were compared to online databases, massbank (massbank.jp) and plantCyc (plantcyc.org). The XCMS package for peak detection in R (cran.r-project.org) was used to align chromatograms, adjust retention times, and group the peaks. For every molecule, a “barcode” peak was chosen to have a unique retention time and mass to charge ratio (m/z) combination. The size of these peaks relative to the internal standard, Quercetin, was used to quantify each molecule in every sample.

## Statistical analysis

The peaks intensities relative to the internal standard were used to capture molecule concentration variation. Standardized intensities were square-root transformed before analysis. Heritability of individual compounds in the three conditions were performed using random intercept models identical to those used to estimate OTU heritability. A fixed “site” effect was added for the field samples. In the greenhouse and sterile conditions, a simple random accession term was used to quantify heritability and estimate accession effects (blups). Those accession effects were used to estimate genetic correlation between specialized metabolites field and greenhouse. We used Pearson’s correlation coefficient and corrected the corresponding p-values for false discovery rate (FDR, N=20).

For the field samples we modeled the relationships between the relative abundances of 19 microbial hubs and the relative intensity of 20 compounds (Extended Data Table 6) using a linear models following:

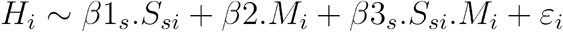

where *H_i_* are the log-ratio transformed counts of one of the 19 microbial hubs used for mapping, and *β1_s_* are the four site effects,*S_si_* is the design matrix assigning sample i to site s, and *β^2^* is the effect of one of the 20 molecules identified in our untargeted screen, is the relative intensity of the molecules measured in sample i. *β3_s_* are site specific regression coefficients (interactions between the site and molecule effects). We fitted 380 models (19 hubs and 20 molecules) and used F-tests to estimate term significance. All *p*-values corresponding to the molecule effect *β^2^* were corrected for False Discovery Rate (N=380).

## Acknowledgements

Many thanks to Mia Holm for her hospitality and wonderful diners after hard work in the field as well as help during harvesting; to Einar Holm for helping with field work and taking photos of harvested plants; to Torbjörn Säll for assistance with sampling and providing greenhouse space in Lund; and finally to the Kleen family, the Öhman family, Nils Jönsson and the Rathckegården farm for allowing us to install our experiments on their land. Thanks to Timothée Flutre and Talia Karasov for helpful discussions on previous versions of the manuscript. Thanks to Man Yu from the Dean lab who helped generate stem images used for seed production estimates and manual seed production estimates. This work was funded by a grant from the National Health Institute (R01 GM 083068) to JB, MN and CD, by a Dropkin Foundation Fellowship to BB and with support from the University of Chicago to JB. BB has received the support of the EU in the framework of the Marie-Curie FP7 COFUND People Programme, through the award of an AgreenSkills/AgreenSkills+ fellowship (under grant agreement n° 267196). Computing resources and storage were provided by: the Center for Research Informatics, funded by the Biological Sciences Division at the University of Chicago with additional funding provided by the Institute for Translational Medicine, CTSA grant number UL1 TR000430 from the National Institutes of Health; the genotoul bioinformatics platform Toulouse Occitanie, France (Bioinfo Genotoul, https://doi.org/10.15454/1.5572369328961167E12); and Bordeaux Bioinformatics Center (CBiB) at the University of Bordeaux, France.

## Author Contributions

BB, DF, SH, JB, MN and CD designed the field trials. BB, DF and SH coordinated fieldwork. BB, DF, EK, FR, AA, MB, SD, TCM, PN, TT, RW took part in fieldwork. HW, with assistance from FH and RF isolated B38, produced the genome sequence and assessed its growth promoting effect in controlled conditions. FR computed the SNP data used for association analysis. BB, PD, MLM, RW produced the microbiota sequence data. PLG, TCM and BB generated and analyzed the metabolomics data. BB and JB conceived of analyses, and BB analyzed the data. BB and JB wrote the paper. MP helped develop the methods for microbiota sequencing. DF, TCM, MP, CD, MN provided comments on the manuscript.

## Repeatability of analysis and data availability

All scripts used to performed the analyses presented in this paper, as well as non-essential but complementary figures, are available in the repository https://forgemia.inra.fr/bbrachi/microbiota_paper.git

The ITS and 16S amplicons and B38 sequence data is available under bioproject <u>PRJNA707473</u>

**Supplementary Information** is linked to the online version of the paper at https://www.nature.com/nature.

## Materials & Correspondence

Reprints and permissions information is available at www.nature.com/reprints

The authors declare no competing financial interests.

## Extended Data

**Extended Data Fig. 1 | Relative frequency of the 10 most frequent OTUs.** Each stacked bar (x-axis) corresponds to a site/year combination. The y-axis gives the proportion of the 10 most frequent OTUs. The colors correspond to the taxonomic assignments of OTUs given in the legend (class / order / family).

**Extended Data Fig. 2 | The effect of host genetic variation on the microbial community targets relatively few OTUs and percolates through hubs**. This figure corresponds to observations in the set of 4 experiments performed in 2012. The same figure is available for the 2013 experiments in Figure 1. **A-D:** Each frame presents the distribution of heritability estimates for individual OTUs in one site. In each frame, the inset graph is a box and whiskers plot contrasting the heritability (y-axis) of bacterial (B) and fungal (F) OTUs. **E-F:** The heritable hubs are represented by large dots, at a distance of 0 (hub). The other OTUs are represented by smaller dots and the x-axis represents their distance to the nearest heritable hub(s) within the sparse covariance networks. The number of heritable hubs detected in each experiment is indicated in the legend. The correlation coefficients presented are Kendall rank correlations calculated for OTUs with a distance to the heritable hub(s) above 0.

**Extended Data Fig. 3 |** Relationship between the mean per site / year combination of the normalized rank abundance of OTUs (x-axis, rank divided by the number of OTUs) in each sample, and heritability (y-axis). Colored points are heritable OTUs and the color and shape indicate the site and year, respectively. Normalized rank abundance of OTUs displays a positive weak but significant relationship with heritability which has an adjusted *r*-squared of 0.04674 (Fstat=205.8, df=4176, p-value: < 2.2e-16).

**Extended Data Fig. 4 | Hubs in microbial networks.** Each frame presents the relationship between degree and betweenness centrality for vertices in the networks computed for each site (SU, SR, NM and NA) and year (2012, 2013). Each dot represents an OTU (fungal or bacterial). The larger and labeled dots correspond to OTUs that have values of betweenness centrality and degree in the 5% tail of both statistics.

**Extended data Fig. 5 | Relationship between prevalence, heritability (A), betweenness (B) and degree (C).** We performed 8 independent experiments, over two years. For each experiment, we defined prevalent OTUs as those detected in over 50% of the plants. In the three panels, the x-axis represents the number of experiments (from 1 to 8) in which an OTU was prevalent, with years distinguished by shape and sites distinguished by color. In A, the y-axis indicates heritability of OTU relative abundance (i.e. variance explained by a random accession effect) estimated within experiments. Colored points represent OTUs with significant heritability. In B and C, the y-axis indicates betweenness and degree of OTU in networks computed for each experiment and colors points are OTUs defined as hubs.

**Extended Data Fig. 6 | Correlation between lifetime seed production (LSP) estimates obtained by counting and measuring siliques (x-axis) versus automated LSP estimates**. A. Row data and Spearman rho rank correlation coefficient. B. Log transformed data and Pearson’s correlation coefficient. In both panels, outliers are indicated in red.

**Extended Data Fig. 7 | Positive correlations among genotype lifetime seed production (LSP) estimates in different experiments.** We measure LSP, a major component of fitness in this autogamous selfing species, in four sites over two years for 200 Swedish accessions. This figure shows the pairwise correlations between accession effects on this fitness component estimated in the eight experiments.

**Extended Data Fig. 8 | Abundant plant specialized metabolites contribute to shaping the relative abundance of microbial hubs. A. Relationships between specialized metabolites and microbial hubs across experiments.** Each bar corresponds to an F-statistic for the effects of the site (grey), the molecule (blue) and the interaction between the two (orange) in a model following the formula HUB ~ Molecule + Site + Molecule *Site (in the form HUB ~ Molecule along the x-axis). The stars associated with each bar indicate the level of significance of the Molecule effect (after FDR correction for 623 tests, only models with *p*-value <0.01 for the molecule effects are shown). Site effects were large for all hubs but the interactions between site and molecule were always small and generally not significant (33 significant in 623 tests without FDR correction; only one significant with FDR correction). **B. Heritability estimates of the molecules** in the field (grey bars) and in the greenhouse (blue bars), and in sterile conditions (orange bars) for each molecule. The vertical segments are 95% confidence intervals obtained with 500 bootstraps for heritability estimates. **C. Genetic correlations for specialized metabolites between accessions grown in the field and in the greenhouse.** Each bar represents a Pearson’s correlation coefficient between field and greenhouse estimates of accession effects (blups) and significance is given by the stars (after FDR correction for 17 tests). Missing bars correspond to molecules with no heritability in the greenhouse and/or the field. B and C share the x-axis labels.

**Extended Data Table 1 | Host variation has a subtle impact on overall community variation**. The first 3 columns indicate the community, site and year for which the analyses were performed. Nh stands for the number of principal coordinate components with significant broad sense heritability estimates (95% confidence intervals not overlapping 0). A total of 10 components were computed for each community/site/year combination. “VE” indicates the total amount of microbial community variation captured by the first 10 components and “he” provides an estimated proportion of total variation explained by the identity of host accessions (over the i heritable components for each site/year combination). The overall host effects reported in the main text reflect the distribution of VE*he in this table.

**Extended Data Table 2 | List of heritable hubs.** Hub OTUs detected in each site and year. H2 is the point heritability estimate for each hub. The columns order, family and genus provide taxonomic assignments.

**Extended Data Table 3 | Hubs are enriched for interkingdom connections (edges).** For each site (first column) and year (second column), the table presents the results from a χ^2^ testing for enrichment in interkingdom edges (third column) when considering all edges, or edges involving at least one hub. B_B, B_F, F_F give the number of edges between 2 bacterial OTUs, a bacterial and a fungal OTU, and 2 fungal OTUs, respectively. The following columns are chi-square values, *p-*values and FDR adjusted *p*-values for 8 tests.

**Extended Data Table 4 | Relationships between host genotype lifetime seed production and influence over microbial hubs.** For each experiment, we computed a multiple linear regression aimed at explaining variation in lifetime seed production among accessions as a function of variation in the effects of accessions on heritable microbial hubs (as well as their squared values indicated by ^“2”^, for example F8 and F8^2^). The table summarizes the results for each site and year, giving the number of accessions used and the adjusted r^2^ for each model after forward/backward model selection. The column “selected terms” indicate the microbial hubs included in the final model, the sign of the effect (-, +) with the significance in the last column (ns: *p*-value ≥ 0.1,.: 0.1 ≥ *p*-value > 0.05, *: 0.05 ≥ *p*-value > 0.01, **: 0.01 ≥ *p*-value > 0.001, ***: *p*-value ≤ 0.001).

**Extended Data Table 5 | Geographical coordinates of Swedish collection sites for live microbial isolates.**

**Extended Data Table 6 | Secondary metabolites detected in this study.** “ID” refers to the identifier assigned to each molecule. “Name” indicates the putative names for the molecules if identified. “Category” describes the type of metabolite: C stands for cyanidin, F stands for flavonoid; GSL stands for glucosinolate; O stands for other. “Base structure” describes the flavonol core of the flavonoids: C stands for Cyanidin, K for Kaempferol and Q for Quercetin. The next eight columns indicate the numbers of different saccharides or the chemical groups that enter in the structure of molecules. “RT” stands for retention time (in second). “mass” indicates the molecular weight: “(obs)” stands for observed and “(exp)” for expected according to the formula.

## Supplementary information

**Supplementary Table 1 |** Natural accessions of *Arabidopsis thaliana* originating from Sweden and grown in 4 sites across Sweden.

**Supplementary Table 2 | Bacterial and Fungal OTUs detected.** The table provides, for the 581 Bacterial OTUs and 704 fungal OTUs, the taxonomic assignations. in addition column “heritable”, “hubs”, “heritable hub” indicate the number of experiment (0 to 8) in which OTUs were significantly influenced by host genotype, a hub in the community and both, respectively. Column “Nexp” indicates the number of experiments in which each OTU was prevalent. “Core microbiota” indicates if the OTU was part of the core microbiota defined in this study (1: yes, 0: no).

**Supplementary Table 3 | QTLs associated with host effects on hubs and our fitness estimate across experiments.** The columns “chromosome”, “start”, and “stop” indicate the genomic coordinates for each QTL. The columns “Nqtl” indicates the number of overlapping associated loci identified by the local score approach which were merged into the QTL. The column “repres” provides a representative SNP for each associated loci aforementioned. Representative SNPs are chosen to have the largest absolute effect on the phenotype for each associated loci. The following column describes which traits display associations with each QTL. For example on line 2, the QTL region overlaps with a loci associated with B41 (value =1) and is an exact match for the loci associated with B99 (value =2). The column “Ntraits” simply counts the number of traits with associations in a QTL region and the column “sizes” is simply the difference between “start” and “stop” and measures QTLs sizes in base pairs.

**Supplementary Table 4 | Biological processes** significantly enriched among genes overlapping with QTLs for microbial hub variation. “trait” simply indicates the trait for which we detected significant enrichment. The columns “name”, “description” and “databases refer to GO terms identification. and “pathway” is the pathway description. “size” refers to the number of genes annotated with the corresponding terms, “setRank” is the setRank statistic characterizing the importance of a gene set, i.e. how much it overlaps with other gene sets, “pSetRank” expresses the probability of observing a gene set with the same setRank value in a random network with the same number of nodes and edges as the observed gene set network. “correctedPValue” is the enrichment *p*-value accounting for overlapping gene sets and “adjustedPValue” is the same probability but adjusted for multiple testing. “enr” and “pv” are the enrichment and associated *p*-value for the method accounting for linkage disequilibrium and non-random distribution of terms along the genome.

**Supplementary Table 5 | Pathways** significantly enriched among genes overlapping with QTLs for microbial hub variation. (See description Supplementary Table 4).

