## Extended data for "Plant genetic effects on microbial hubs impact fitness across field trials"

**for**

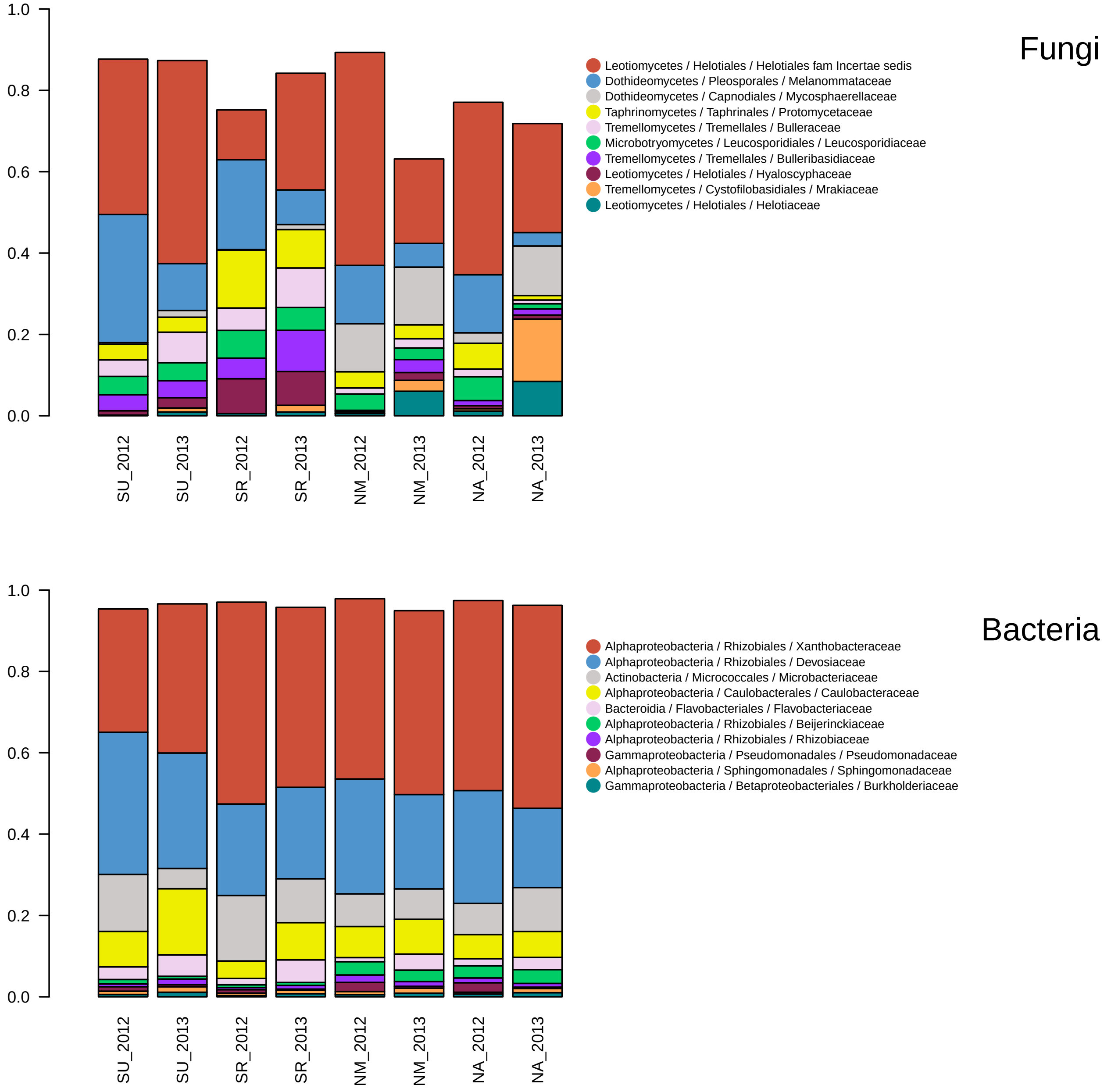


**Extended Data Fig. 1 | Relative frequency of the 10 most frequent OTUs.** Each stacked bar (x-axis) corresponds to a site/year combination. The y-axis gives the proportion of the 10 most frequent OTUs. The colors correspond to the taxonomic assignments of OTUs given in the legend (class / order / family).

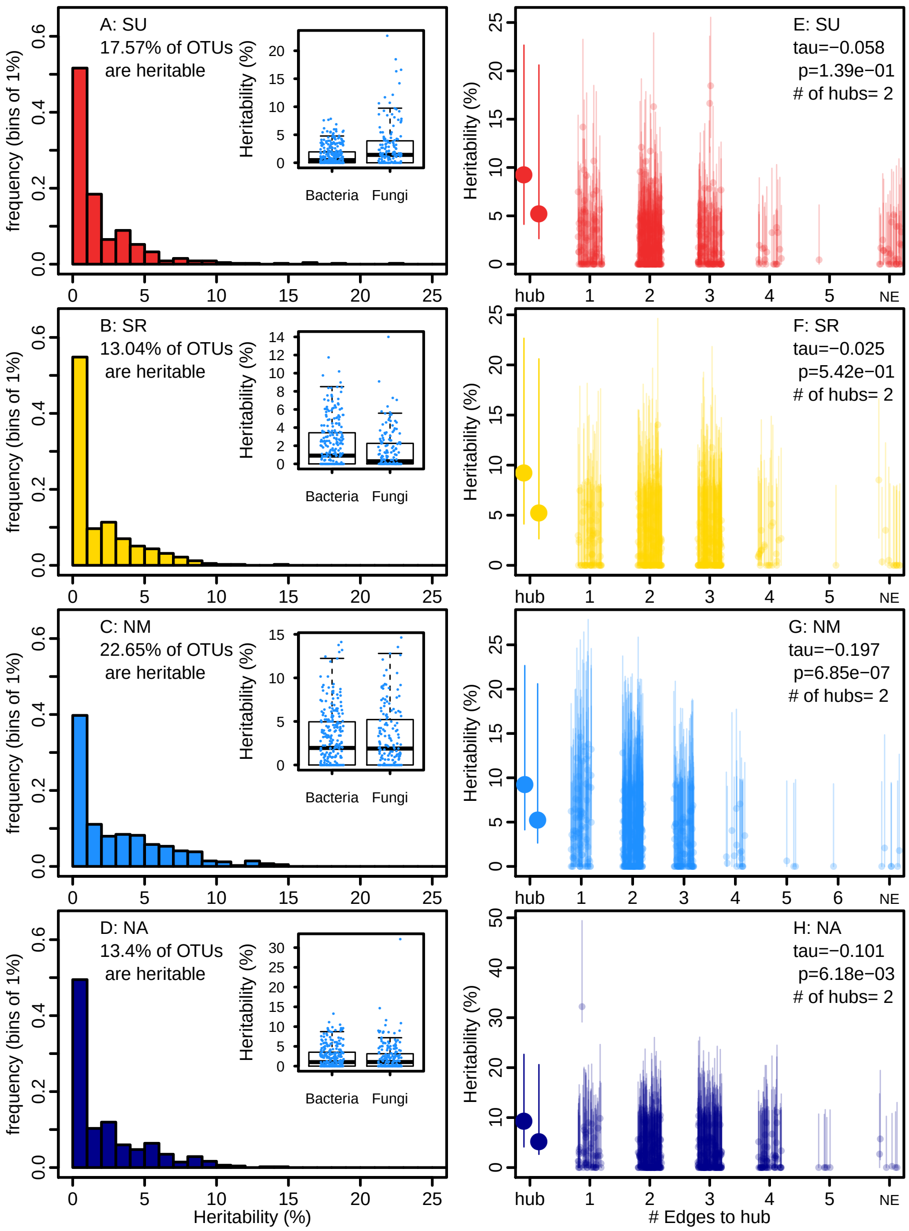


**Extended Data Fig. 2 | The effect of host genetic variation on the microbial community targets relatively few OTUs and percolates through hubs**. This figure corresponds to observations in the set of 4 experiments performed in 2012. The same figure is available for the 2013 experiments in Figure 1. **A-D:** Each frame presents the distribution of heritability estimates for individual OTUs in one site. In each frame, the inset graph is a box and whiskers plot contrasting the heritability (y-axis) of bacterial (B) and fungal (F) OTUs. **E-F:** The heritable hubs are represented by large dots, at a distance of 0 (hub). The other OTUs are represented by smaller dots and the x-axis represents their distance to the nearest heritable hub(s) within the sparse covariance networks. The number of heritable hubs detected in each experiment is indicated in the legend. The correlation coefficients presented are Kendall rank correlations calculated for OTUs with a distance to the heritable hub(s) above 0.

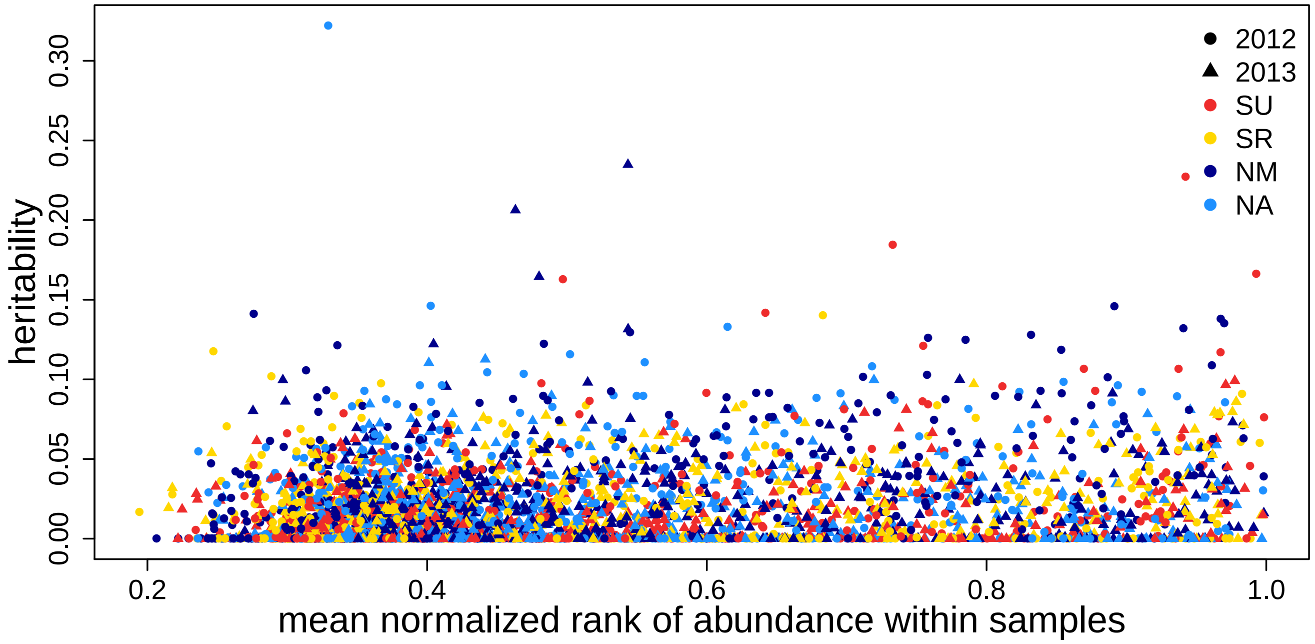


**Extended Data Fig. 3 |** Relationship between the mean per site / year combination of the normalized rank abundance of OTUs (x-axis, rank divided by the number of OTUs) in each sample, and heritability (y-axis). Colored points are heritable OTUs and the color and shape indicate the site and year, respectively. Normalized rank abundance of OTUs displays a positive weak but significant relationship with heritability which has an adjusted *r*-squared of 0.04674 (Fstat=205.8, df=4176, p-value: < 2.2e-16).

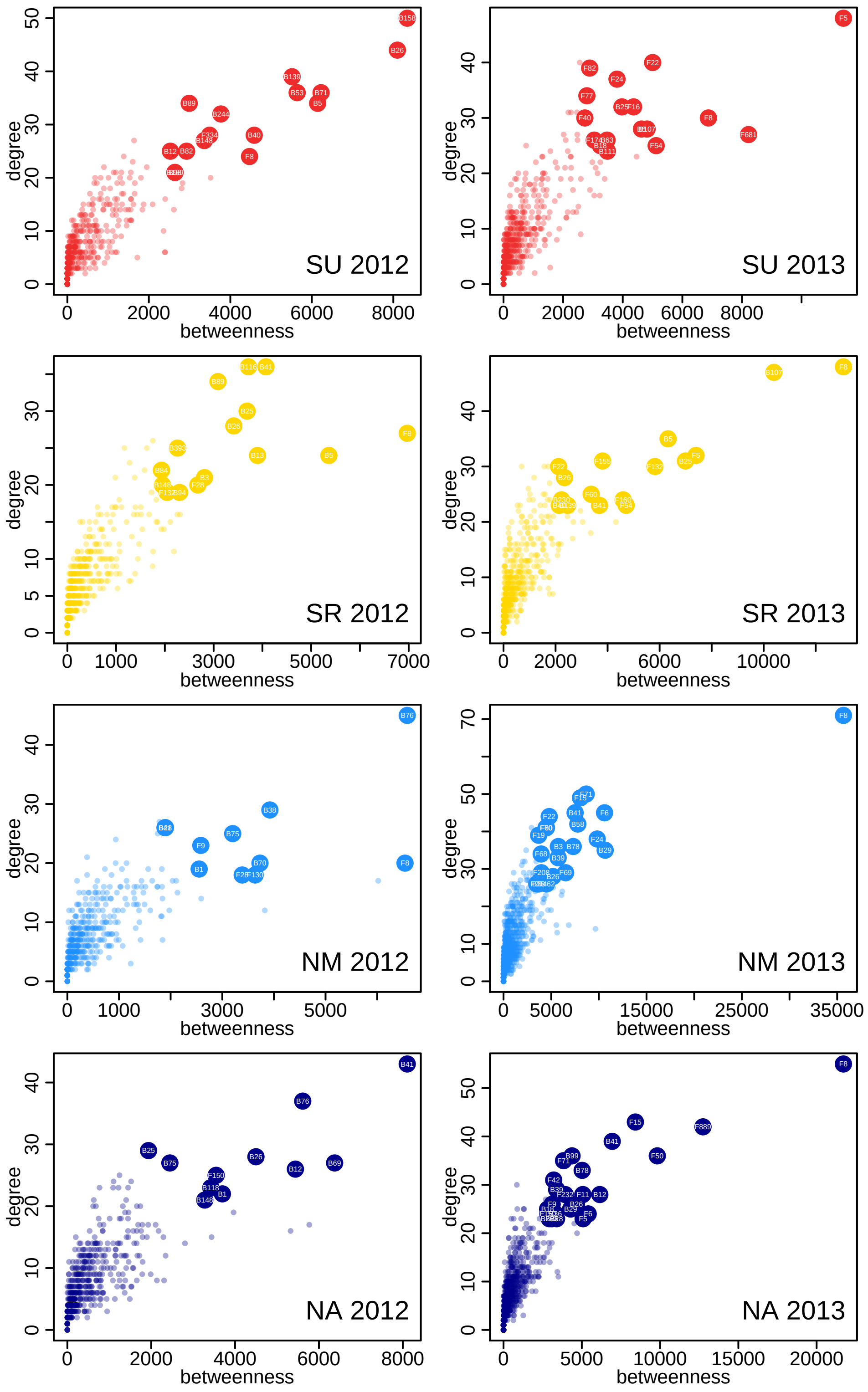


**Extended Data Fig. 4 | Hubs in microbial networks.** Each frame presents the relationship between degree and betweenness centrality for vertices in the networks computed for each site (SU, SR, NM and NA) and year (2012, 2013). Each dot represents an OTU (fungal or bacterial). The larger and labeled dots correspond to OTUs that have values of betweenness centrality and degree in the 5% tail of both statistics.

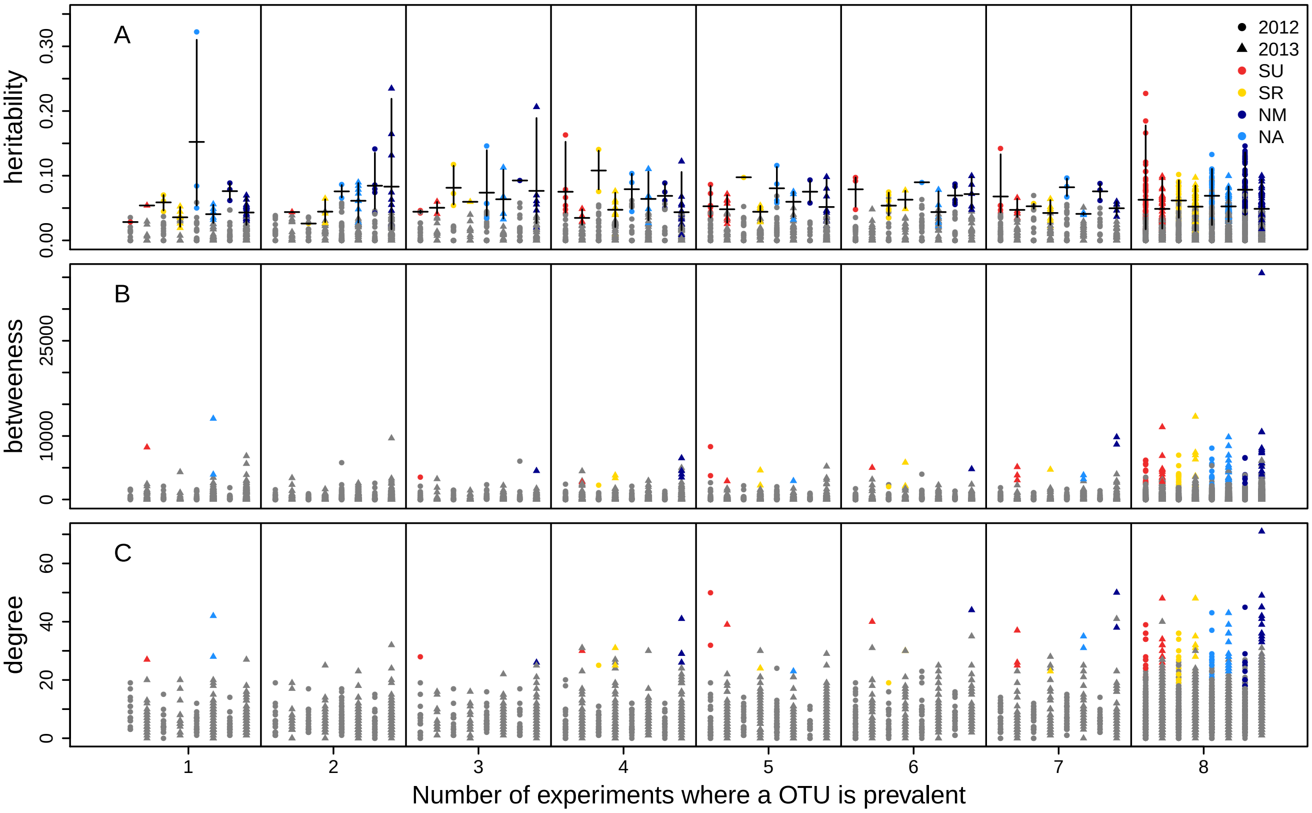


**Extended data Fig. 5 | Relationship between prevalence, heritability (A) , betweenness (B) and degree (C).** We performed 8 independent experiments, over two years. For each experiment, we defined prevalent OTUs as those detected in over 50% of the plants. In the three panels, the x-axis represents the number of experiments (from 1 to 8) in which an OTU was prevalent, with years distinguished by shape and sites distinguished by color. In A, the y-axis indicates heritability of OTU relative abundance (i.e. variance explained by a random accession effect) estimated within experiments. Colored points represent OTUs with significant heritability. In B and C, the y-axis indicates betweenness and degree of OTU in networks computed for each experiment and colors points are OTUs defined as hubs.

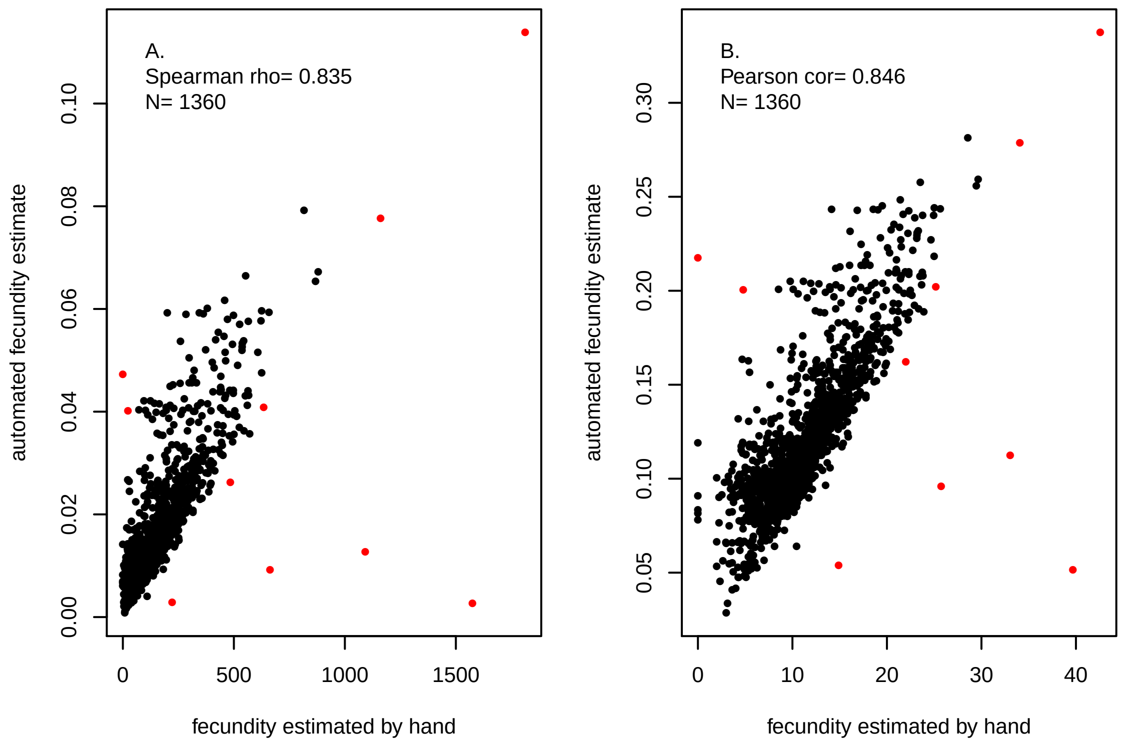


**Extended Data Fig. 6 | Correlation between lifetime seed production (LSP) estimates obtained by counting and measuring siliques (x-axis) versus automated LSP estimates**. A. Row data and Spearman rho rank correlation coefficient. B. Log transformed data and Pearson's correlation coefficient. In both panels, outliers are indicated in red.

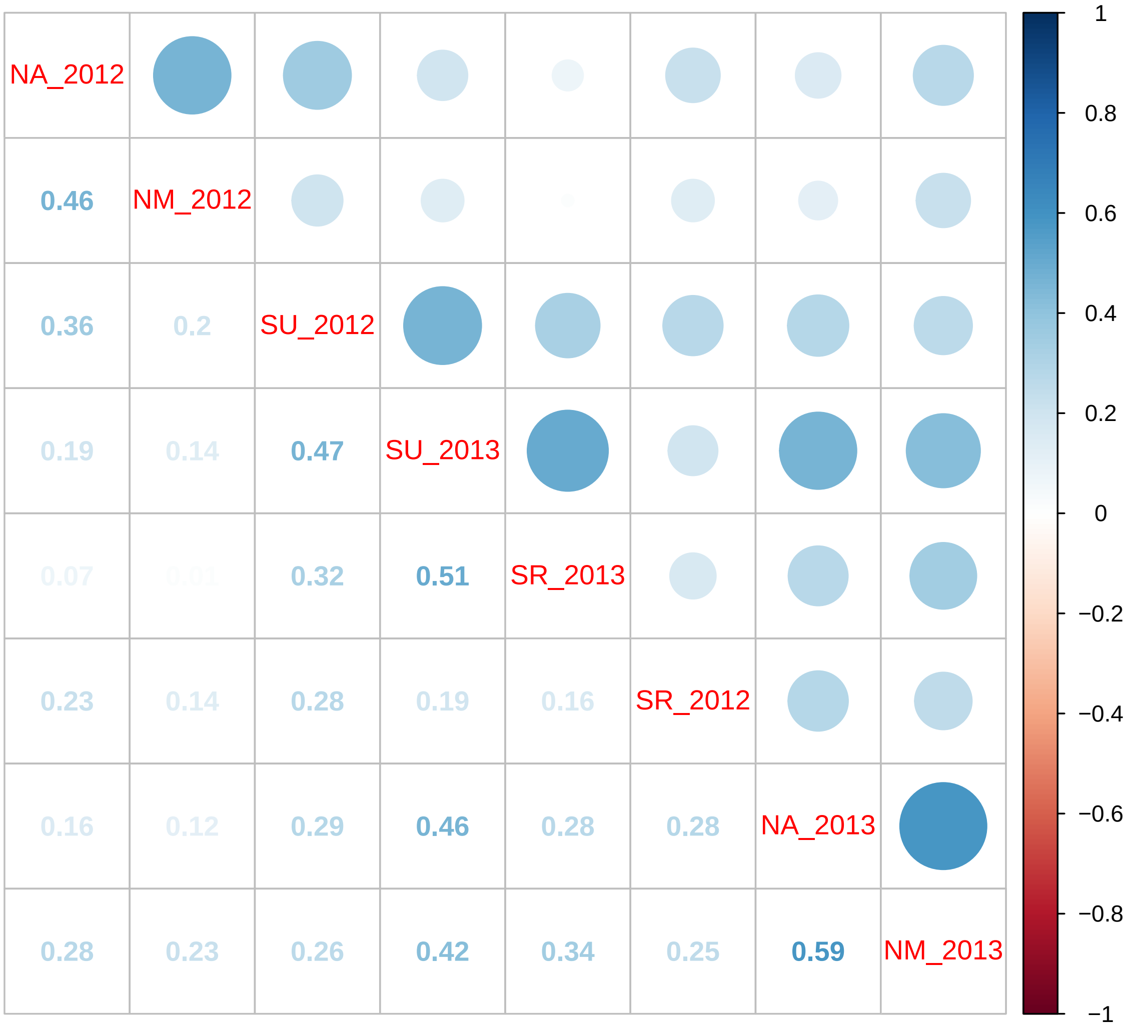


**Extended Data Fig. 7 | Positive correlations among genotype lifetime seed production (LSP) estimates in different experiments.** We measure LSP, a major component of fitness in this autogamous selfing species, in four sites over two years for 200 Swedish accessions. This figure shows the pairwise correlations between accession effects on this fitness component estimated in the eight experiments.

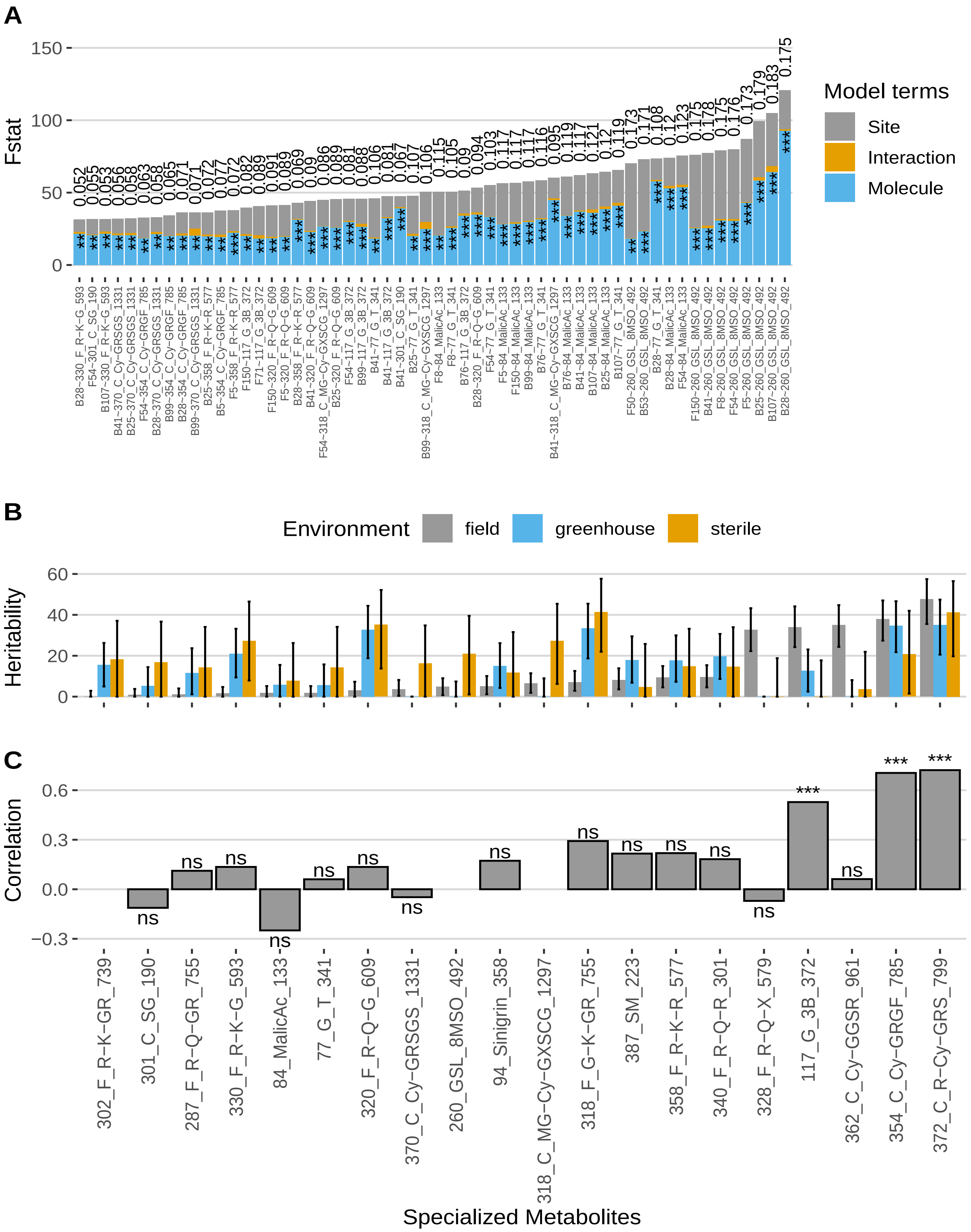

A
Fstat
150
100
50
0
Model terms
Site
Interaction
Molecule
B28~330_F_R~K~G_593
F54~301_C_SG_190
B107~330_F_R~K~G_593
B41~370_C_Cy~GRSGS_1331
B25~370_C_Cy~GRSGS_1331
F54~354_C_Cy~GRGF_785
B28~370_C_Cy~GRSGS_1331
B99~354_C_Cy~GRGF_785
B28~354_C_Cy~GRGF_785
B99~370_C_Cy~GRSGS_1331
B25~358_F_R~K~R_577
B5~354_C_Cy~GRGF_785
F5~358_F_R~K~R_577
F150~117_G_3B_372
F71~117_G_3B_372
F150~320_F_R~Q~G_609
F5~320_F_R~Q~G_609
B28~358_F_R~K~R_577
B41~320_F_R~Q~G_609
F54~318_C_MG~Cy~GXSCG_1297
B25~320_F_R~Q~G_609
F54~117_G_3B_372
B99~117_G_3B_372
B41~77_G_T_341
B41~117_G_3B_372
B41~301_C_SG_190
B25~77_G_T_341
B99~318_C_MG~Cy~GXSCG_1297
F8~84_MalicAc_133
F8~77_G_T_341
B76~117_G_3B_372
B28~320_F_R~Q~G_609
F54~77_G_T_341
F5~84_MalicAc_133
F150~84_MalicAc_133
B99~84_MalicAc_133
B76~77_G_T_341
B41~318_C_MG~Cy~GXSCG_1297
B76~84_MalicAc_133
B41~84_MalicAc_133
B107~84_MalicAc_133
B25~84_MalicAc_133
B107~77_G_T_341
F50~260_GSL_8MSO_492
B53~260_GSL_8MSO_492
B28~77_G_T_341
B28~84_MalicAc_133
F54~84_MalicAc_133
F150~260_GSL_8MSO_492
B41~260_GSL_8MSO_492
F8~260_GSL_8MSO_492
F54~260_GSL_8MSO_492
F5~260_GSL_8MSO_492
B25~260_GSL_8MSO_492
B107~260_GSL_8MSO_492
B28~260_GSL_8MSO_492
B
Environment
field
greenhouse
sterile
Heritability
60
40
20
0
C
Correlation
0.6
0.3
0.0
−0.3
ns
***
302_F_R−K−GR_739
301_C_SG_190
287_F_R−Q−GR_755
330_F_R−K−G_593
84_MalicAc_133
77_G_T_341
320_F_R−Q−G_609
370_C_Cy−GRSGS_1331
260_GSL_8MSO_492
94_Sinigrin_358
318_C_MG−Cy−GXSCG_1297
318_F_G−K−GR_755
387_SM_223
358_F_R−K−R_577
340_F_R−Q−R_301
328_F_R−Q−X_579
117_G_3B_372
362_C_Cy−GGSR_961
354_C_Cy−GRGF_785
372_C_R−Cy−GRS_799
Specialized Metabolites

**Extended Data Figure 8 | Abundant plant specialized metabolites contribute to shaping the relative abundance of microbial hubs. A. Relationships between specialized metabolites and microbial hubs across experiments.** Each bar corresponds to an F-statistic for the effects of the site (grey), the molecule (blue) and the interaction between the two (orange) in a model following the formula HUB ~ Molecule+ Site + Molecule *Site (in the form HUB ~ Molecule along the x-axis). The stars associated with each bar indicate the level of significance of the Molecule effect (after FDR correction for 623 tests, only models with $p$-value <0.01 for the molecule effects are shown). Site effects were large for all hubs but the interactions between site and molecule were always small and generally not significant (33 significant in 623 tests without FDR correction; none significant with FDR correction). **B. Heritability estimates of the molecules** in the field (grey bars) and in the greenhouse (blue bars), and in sterile conditions (orange bars) for each molecule. The vertical segments are 95% confidence intervals obtained with 500 bootstraps for heritability estimates. **C. Genetic correlations for specialized metabolites between accessions grown in the field and in the greenhouse**. Each bar represents a Pearson's correlation coefficient between field and greenhouse estimates of accession effects (blups) and significance is given by the stars (after FDR correction for 17 tests). Missing bars correspond to molecules with no heritability in the greenhouse and/or the field. B and C share the x-axis labels.

**Extended Data Table 1 | Host variation has subtle impact on overall community variation.**

| community | site | year | Nh | VE | he |
|---|---|---|---|---|---|
| Fungal | SU | 2012 | 9 | 58.72 | 5.22 |
| | | 2013 | 9 | 56.25 | 0.81 |
| | SR | 2012 | 10 | 47.99 | 2.28 |
| | | 2013 | 10 | 49.50 | 3.00 |
| | NM | 2012 | 10 | 60.07 | 2.80 |
| | | 2013 | 8 | 42.54 | 1.14 |
| | NA | 2012 | 10 | 61.58 | 1.78 |
| | | 2013 | 10 | 54.64 | 0.66 |
| Bacterial | SU | 2012 | 7 | 49.24 | 3.49 |
| | | 2013 | 8 | 41.47 | 1.45 |
| | SR | 2012 | 10 | 57.18 | 1.18 |
| | | 2013 | 10 | 45.90 | 1.99 |
| | NM | 2012 | 10 | 57.68 | 3.80 |
| | | 2013 | 9 | 45.84 | 2.45 |
| | NA | 2012 | 9 | 55.19 | 1.28 |
| | | 2013 | 10 | 51.78 | 1.64 |

The first 3 columns indicate the community, site and year for which the analyses were performed. Nh stands for the number of principal coordinate components with significant broad sense heritability estimates (95% confidence intervals not overlapping 0). A total of 10 components were computed for each community/site/year combination. "VE" indicates the total amount of microbial community variation captured by the first 10 components and "he" provides an estimated proportion of total variation explained by the identity of host accessions (over the i heritable components for each site/year combination). The overall host effects reported in the main text reflect the distribution of VE*he in this table.

**Extended Data Table 2 | List of heritable hubs.**

| OTU | exp | year | h2 | order | family | genus | species |
|---|---|---|---|---|---|---|---|
| B107 | SU | 2013 | 0.0792 | Betaproteobacteriales | Burkholderiaceae | arcticum group | uncultured bacterium |
| B12 | NA | 2013 | 0.0809 | Sphingomonadales | Sphingomonadaceae | Sphingomonas | Sphingomonas aquatilis |
| B13 | SR | 2012 | 0.0628 | Betaproteobacteriales | Burkholderiaceae | NA | NA |
| B25 | SU | 2013 | 0.0685 | | | | |
| B25 | SR | 2013 | 0.0348 | | | | |
| B26 | SR | 2013 | 0.0793 | Rhizobiales | Beijerinckiaceae | Methylobacterium | uncultured bacterium |
| B26 | NA | 2013 | 0.0501 | | | | |
| B26 | NA | 2012 | 0.0923 | | | | |
| B28 | NM | 2012 | 0.1014 | Betaproteobacteriales | Burkholderiaceae | Polaromonas | NA |
| B38 | NM | 2012 | 0.0736 | Caulobacterales | Caulobacteraceae | Brevundimonas | Ambiguous taxa |
| B41 | SR | 2013 | 0.0111 | Betaproteobacteriales | Burkholderiaceae | NA | NA |
| B41 | SR | 2012 | 0.0713 | | | | |
| B5 | SU | 2013 | 0.0315 | Betaproteobacteriales | Burkholderiaceae | Variovorax | NA |
| B53 | SU | 2012 | 0.0177 | Sphingomonadales | Sphingomonadaceae | Sphingomonas | uncultured soil bacterium |
| B76 | NA | 2012 | 0.0522 | Corynebacteriales | Nocardiaceae | Nocardia | actinobacterium P23 |
| B99 | NA | 2013 | 0.0996 | Betaproteobacteriales | Burkholderiaceae | Rhizobacter | Ambiguous taxa |
| F130 | NM | 2012 | 0.0601 | NA | NA | NA | NA |
| F150 | NA | 2013 | 0.0611 | Taphrinales | Taphrinaceae | Taphrina | Taphrina tormentillae |
| F160 | SR | 2013 | 0.0479 | Pleosporales | Phaeosphaeriaceae | Phaeosphaeria | Phaeosphaeria caricicola |
| F334 | SU | 2012 | 0.0464 | Pleosporales | Pleosporaceae | Alternaria | Alternaria lolii |
| F5 | SU | 2013 | 0.0628 | Capnodiales | Mycosphaerellaceae | Mycosphaerella | Mycosphaerella tassiana |
| F5 | SR | 2013 | 0.0796 | | | | |
| F50 | NA | 2013 | 0.0292 | NA | NA | NA | NA |
| F54 | SR | 2013 | 0.0458 | Sporidiobolales | Sporidiobolaceae | Sporobolomyces | Sporobolomyces roseus |
| F60 | SR | 2013 | 0.0588 | Leucosporidiales | Leucosporidiaceae | Leucosporidium | Leucosporidium yakuticum |
| F60 | NM | 2013 | 0.0451 | | | | |
| F69 | NM | 2013 | 0.0083 | NA | NA | NA | NA |
| F71 | NM | 2013 | 0.0463 | Helotiales | Helotiaceae | Tetracladium | Tetracladium marchalianum |
| F71 | NA | 2013 | 0.0393 | | | | |
| F8 | SU | 2012 | 0.2271 | Taphrinales | Protomycetaceae | Protomyces | Protomyces inouyei |
| F8 | NM | 2012 | 0.1260 | | | | |
| F8 | SU | 2013 | 0.0966 | | | | |
| F8 | SR | 2013 | 0.0716 | | | | |
| F8 | NM | 2013 | 0.0731 | | | | |

Hub OTUs detected in each site and year. H2 is the point heritability estimate for each hub. The columns order, family and genus provide taxonomic assignments.

**Extended Data Table 3 | Hubs are enriched for interkingdom connections (edges).**

| site | year | edges | B_B | B_F | F_F | chisq | pval | adjpval |
|---|---|---|---|---|---|---|---|---|
| SU | 2012 | all | 1148 | 136 | 555 | 106.43 | <2e-05 | 1.60E-04 |
| | | withhubs | 369 | 61 | 34 | | | |
| SU | 2013 | all | 1173 | 249 | 978 | 73.63 | <2e-05 | 1.60E-04 |
| | | withhubs | 143 | 86 | 276 | | | |
| SR | 2012 | all | 980 | 136 | 483 | 41.30 | <2e-05 | 1.60E-04 |
| | | withhubs | 276 | 40 | 50 | | | |
| SR | 2013 | all | 1228 | 249 | 991 | 32.46 | <2e-05 | 1.60E-04 |
| | | withhubs | 198 | 88 | 169 | | | |
| NA | 2012 | all | 1218 | 158 | 638 | 70.79 | <2e-05 | 1.60E-04 |
| | | withhubs | 240 | 29 | 23 | | | |
| NA | 2013 | all | 1433 | 288 | 1184 | 21.03 | 6e-05 | 4.80E-04 |
| | | withhubs | 290 | 105 | 294 | | | |
| NM | 2012 | all | 1077 | 117 | 491 | 7.66 | 0.022 | 1.79E-01 |
| | | withhubs | 174 | 27 | 58 | | | |
| NM | 2013 | all | 2049 | 358 | 1737 | 84.21 | <2e-05 | 1.60E-04 |
| | | withhubs | 273 | 130 | 406 | | | |

For each site (first column) and year (second column), the table presents the results from a $\chi 2$ testing for enrichment in interkingdom edges (third column) when considering all edges, or edges involving at least one hub. B_B, B_F, F_F give the number of edges between 2 bacterial OTUs, a bacterial and a fungal OTU, and 2 fungal OTUs, respectively. The following columns are chi-square values, *p*-values and FDR adjusted *p*-values for 8 tests.

**Extended Data Table 4 | Relationships between host genotype lifetime seed production and influence over microbial hubs.**

| year | site | terms | estimate | std error | t.value | p.value | significance |
|---|---|---|---|---|---|---|---|
| 2012 | SU | (Intercept) | -0.001 | 0.001 | -1.079 | 2.82E-01 | ns |
| | | B53 | 0.055 | 0.015 | 3.595 | 4.12E-04 | *** |
| | | F8 | 0.010 | 0.002 | 6.529 | 5.75E-10 | *** |
| | | $F8^2$ | 0.004 | 0.002 | 2.211 | 2.82E-02 | * |
| 2013 | SU | (Intercept) | 0.000 | 0.001 | 0.310 | 7.57E-01 | ns |
| | | F8 | 0.018 | 0.004 | 4.845 | 2.56E-06 | *** |
| 2012 | SR | (Intercept) | 0.000 | 0.001 | -0.162 | 8.71E-01 | ns |
| | | B13 | 0.025 | 0.009 | 2.740 | 6.79E-03 | ** |
| | | $B13^2$ | 0.123 | 0.078 | 1.572 | 1.18E-01 | ns |
| 2013 | SR | (Intercept) | 0.000 | 0.001 | -0.004 | 9.97E-01 | ns |
| | | B25 | 0.013 | 0.009 | 1.492 | 1.37E-01 | ns |
| | | B26 | 0.010 | 0.004 | 2.378 | 1.84E-02 | * |
| | | B41 | -0.054 | 0.026 | -2.119 | 3.54E-02 | * |
| | | F160 | 0.006 | 0.004 | 1.409 | 1.60E-01 | ns |
| | | F5 | -0.014 | 0.004 | -3.632 | 3.64E-04 | *** |
| | | F60 | 0.010 | 0.004 | 2.582 | 1.06E-02 | * |
| | | F8 | 0.013 | 0.004 | 3.071 | 2.45E-03 | ** |
| | | $B41^2$ | -1.308 | 0.715 | -1.828 | 6.91E-02 | . |
| | | $F8^2$ | 0.033 | 0.016 | 2.102 | 3.69E-02 | * |
| 2013 | NA | (Intercept) | -0.001 | 0.001 | -1.004 | 3.16E-01 | ns |
| | | B26 | 0.013 | 0.008 | 1.749 | 8.19E-02 | . |
| | | $F150^2$ | 0.048 | 0.024 | 1.990 | 4.80E-02 | * |
| 2012 | NM | (Intercept) | 0.000 | 0.001 | 0.046 | 9.63E-01 | ns |
| | | $B28^2$ | 0.055 | 0.032 | 1.746 | 8.27E-02 | . |
| 2013 | NM | (Intercept) | 0.000 | 0.001 | 0.475 | 6.35E-01 | ns |
| | | F60 | -0.028 | 0.008 | -3.723 | 2.58E-04 | *** |
| | | F69 | -0.077 | 0.034 | -2.281 | 2.36E-02 | * |
| | | F8 | 0.012 | 0.006 | 2.063 | 4.05E-02 | * |

For each experiment, we computed a multiple linear regression aimed at explaining variation in lifetime seed production among accessions as a function of variation in the effects of accessions on heritable microbial hubs (as well as their squared values indicated by "$^2$", for example F8 and $F8^2$) . The table summarizes the results for each site and year, giving the number of accessions used and the adjusted $r^2$ for each model after forward/backward model selection. The column "selected terms" indicate the microbial hubs included in the final model, the sign of the effect (-, +) with the significance in the last column (ns: $p$-value $\geq 0.1$, . : $0.1 \geq p$-value $> 0.05$, *: $0.05 \geq p$-value $> 0.01$, **: $0.01 \geq p$-value $> 0.001$, *** : $p$-value $\leq 0.001$).

**Extended Data Table 5 | Geographical coordinates of Swedish collection sites for live microbial isolates.**

| Collection | Patch | Site Lat (N) | Site Long (E) | Sample ID | Date |
|---|---|---|---|---|---|
| **Adal (NA)** | Patch #1: 1-10 | 62.86216 | 18.33597 | A | 5-May-17 |
| **Varhallarna (near SR)** | Patch #1: 1-5 | 55.58 | 14.334 | Var | 11-Apr-17 |
| **Varhallarna (near SR)** | Patch #2: 6-10 | 55.58 | 14.334 | Var | 11-Apr-17 |
| **Ullstorp (near SU)** | Patch #1: 1-5 | 56.0648 | 13.9707 | Ull | 10-Apr-17 |
| **Ullstorp (near SU)** | Patch #2: 6-10 | 56.0648 | 13.9707 | Ull | 10-Apr-17 |
| **Tjor (inland) (near SR)** | Patch #1: 1-10 | 58.041 | 11.683 | TJ2 | 12-Apr-17 |
| **Tjor (beach) (near SR)** | Patch #1: 1-5 | 58.041 | 11.683 | Tjor | 12-Apr-17 |
| **Tjor (beach) (near SR)** | Patch #2: 6-8 | 58.041 | 11.683 | Tjor | 12-Apr-17 |
| **Tjor (beach) (near SR)** | Patch #2: 9-10 | 58.041 | 11.683 | Tjor | 12-Apr-17 |

**Extended Data Table 6: Secondary metabolites detected in this study.** “ID” refers to the identifier assigned to each molecule. “Name” indicates the putative names for the molecules if identified. “Category” describes the type of metabolite: C stands for cyanidin, F stands for flavonoid; GSL stands for glucosinolate; O stands for other. “Base structure” describes the flavonol core of the flavonoids: C stands for Cyanidin, K for Kaempferol and Q for Quercetin. The next eight columns indicate the numbers of different saccharides or the chemical groups that enter in the structure of molecules. “RT” stands for retention time (in second). “mass” indicates the molecular weight: “(obs)” stands for observed and “(exp)” for expected according to the formula.

| ID* | Name | Category | Base structure | Glucose | Rhamnose | Xylose | Galactose | Sinapoyl | Malonyl | Coumaroyl | Feruloyl | RT | Mass (obs) | Mass (exp) |
|---|---|---|---|---|---|---|---|---|---|---|---|---|---|---|
| 318_C_MG-**Cy**-GXSCG_1297 | | C | Cy | 2 | | 1 | | 1 | 1 | 1 | | 318 | 1341.3315 | 1341.3363 |
| 354_C_**Cy**-GRGF_785 | | C | Cy | 1 | 1 | | 1 | | | | 1 | 354 | 931.248 | 931.25136 |
| 372_C_R-**Cy**-GRS_799 | | C | Cy | 1 | 2 | | | 1 | | | | 372 | 1052.22 | |
| 362_C_**Cy**-GGSR_961 | | C | Cy | 1 | 1 | | 1 | 1 | | | | 362 | 961.2628 | 961.2608 |
| 370_C_**Cy**-GRSGS_1331 | | C | Cy | 2 | 1 | | | 2 | | | | 370 | 1331.353 | |
| 302_F_R-**K**-GR_739 | | F | K | 1 | 2 | | | | | | | 302 | 739.2091 | 739.2091 |
| 318_F_G-**K**-GR_755 | | F | K | 2 | 1 | | | | | | | 318 | 755.2027 | 755.204 |
| 330_F_R-**K**-G_593 | | F | K | 1 | 1 | | | | | | | 330 | 593.15 | 593.1512 |
| 358_F_R-**K**-R_577 | Kaempferitrin | F | K | | 2 | | | | | | | 358 | 577.1573 | 577.1563 |
| 287_F_R-**Q**-GR_755 | | F | Q | 1 | 2 | | | | | | | 287 | 755.2055 | 755.204 |
| 320_F_R-**Q**-G_609 | | F | Q | 1 | 1 | | | | | | | 320 | 609.1479 | 609.1461 |
| 328_F_R-**Q**-X_579 | | F | Q | | 1 | 1 | | | | | | 328 | 579.1359 | 579.1355 |
| 340_F_R-**Q**-R_301 | | F | Q | | 2 | | | | | | | 340 | 593.1536 | 593.1512 |
| 260_GSL_8MSO_492 | 8-methylsulfinyloctyl | GSL | | 1 | | | | | | | | 260 | 492.1039 | 492.1037 |
| 94_Sinigrin_358 | Sinigrin | GSL | | 1 | | | | | | | | 94 | 358.02 | 358.02 |
| 117_G_3B_372 | 3-butenyl | GSL | | 1 | | | | | | | | 117 | 372.0442 | 372.0428 |
| 301_C_SG_190 | Glucopyranosyl sinapate | O | | 1 | | | | 1 | | | | 301 | 385.1131 | 385.114 |
| 387_SM_223 | Sinapoyl malate | O | | | | | | 1 | 1 | | | 387 | 339.07 | 338.26 |
| 84_MalicAc_133 | Malic acid | O | | | | | | | 1 | | | 84 | 133.03 | 134.08 |
| 77_G_T_341 | Trehalose | O | | 2 | | | | | | | | 77 | 341.1119 | 342.29 |

***Naming convention:** For flavonoids, molecule names in the ID column are formed as follows: RT_Category_CODE_MZ where RT is the retention time observed in seconds, Category can be Cyanidin, or F for flavonols, and MZ is the m/z for the ion which served as a diagnostic tag for the molecule. CODE is build as follows: Bold letters (Cy, K or Q) separated from other letters by dashes refer to the base structure of the molecule. Letters before the core refer to components on #5 carbon for cyanidins and #7 carbon for K or Q; and letters after the core refer to components on the #3 flavonol carbon.
